# Resolving the conflict between associative overdominance and background selection

**DOI:** 10.1101/042390

**Authors:** Lei Zhao, Brian Charlesworth

## Abstract

In small populations, genetic linkage between a polymorphic neutral locus and loci subject to selection, either against partially recessive mutations or in favor of heterozygotes, may result in an apparent selective advantage to heterozygotes at the neutral locus (associative overdominance), and a retardation of the rate of loss of variability by genetic drift at this locus. In large populations, selection against deleterious mutations has previously been shown to reduce variability at linked neutral loci (background selection). We describe analytical, numerical and simulation studies that shed light on the conditions under which retardation versus acceleration of loss of variability occurs at a neutral locus linked to a locus under selection. We consider a finite, randomly mating population initiated from an infinite population in equilibrium at a locus under selection, with no linkage disequilibrium. With mutation and selection, retardation only occurs when *S*, the product of twice the effective population size and the selection coefficient, is of order one. With *S* ≫ 1, background selection always causes an acceleration of loss of variability. Apparent heterozygote advantage at the neutral locus is, however, always observed when mutations are partially recessive, even if there is an accelerated rate of loss of variability. With heterozygote advantage at the selected locus, there is nearly always a retardation of loss of variability. The results shed light on experiments on the loss of variability at marker loci in laboratory populations, and on the results of computer simulations of the effects of multiple selected loci on neutral variability.

---

There has recently been much interest in the effects of selection at one locus on patterns of evolution and variation at linked neutral or nearly-neutral loci, with mounting evidence for such effects from surveys of genome-wide patterns of molecular variability and evolution (Cutter and Payseur 2013; Neher 2013; Charlesworth and Campos 2014). Attention has been especially directed at the possibility of enhanced neutral variability at nucleotide sites that are closely linked to sites under long-term balancing selection (Charlesworth 2006; Gao *et al*. 2015), and at the reduction in variability caused by the hitchhiking effects of directional selection, involving either positive selection (selective sweeps) (Maynard Smith and Haigh 1974) or negative selection (background selection) (Charlesworth *et al*. 1993). There seems little doubt that linkage effects play an important role in shaping patterns of variability across the genome (Cutter and Payseur 2013; Charlesworth and Campos 2014).

Recent studies have, however, made little or no reference to the classical work on associative overdominance (AOD), which dates back to over 40 years ago Following a proposal by Frydenberg (1963), who coined the term, it was shown by Sved (1968, 1971, 1972) and by Ohta and Kimura (Ohta and Kimura 1970; Ohta 1971, 1973) that linkage disequilibrium (LD) between a polymorphic neutral locus and a locus subject either to selection in favour of heterozygotes, or to selection against recessive or partially recessive mutant deleterious alleles, could result in apparent heterozygote advantage at the neutral locus. This is because such LD, generated by genetic drift in a randomly mating finite population, leads to an association between homozygosity at the two loci (Haldane 1949). If homozygosity at the selected locus results in reduced fitness, homozygotes at the neutral locus will also suffer reduced fitness. A heuristic treatment of this effect is given by Charlesworth and Charlesworth (2010, pp. 396-7, 403-4).

This effect of LD in a randomly mating population should be distinguished from the effect of identity disequilibrium (ID) in a population with a mixture of random mating and matings between close relatives (Haldane 1949; Cockerham and Weir 1968), as exemplified by species that reproduce by a mixture of outcrossing and self-fertilization. Here, AOD can arise because of variation among individuals in their inbreeding coefficients, which causes correlations among loci in their levels of homozygosity even if they are unlinked (Haldane 1949; Cockerham and Weir 1968). Theoretical models show that this type of AOD does not require either finite population size or LD among loci (Ohta and Cockerham 1974; Charlesworth 1991). In a finite randomly mating population, however, LD and ID are formally equivalent (Weir *et al*. 1980; Bierne *et al*. 2000).

Associations between heterozygosity at putatively neutral molecular marker loci and higher values of fitness components have frequently been found in natural populations of many different types of organism, leading to a debate as to whether AOD due to LD with random mating or to ID caused by variation in inbreeding levels is the main cause of these associations (David 1998; Hansson and Westerberg 2002). Current evidence appears to favor the latter model (Szulkin *et al*. 2010; Hoffman 2014), especially because the LD model seems to require a relatively small effective population size to produce substantial effects, unless linkage is very tight (Ohta 1971, 1973).

It has also been reported that the level of variability in both quantitative traits and marker loci in laboratory populations maintained with known effective population sizes can sometimes decline less rapidly over time than is predicted by the standard neutral model (e.g., Rumball *et al*. 1994; Gilligan *et al*. 2005; Latter 1998). In addition, laboratory populations and populations of domesticated animals and plants often have levels of quantitative trait variability that are surprisingly high, given their low effective population sizes (Johnson and Barton 2005; Hill 2010). Sved and Ohta proposed that AOD caused by randomly generated LD leads to a retardation in the loss of variability at neutral loci linked to loci under selection; Ohta suggested that this effect could be substantial when the effects of partially recessive mutations distributed over a whole chromosome are considered, provided that the population size is sufficiently small (Ohta 1971, 1973). Computer simulations of multi-locus systems have shown that a retardation of the rate of loss of neutral variability can indeed occur in small populations (Latter 1998; Pamilo and Palsson 1998, 1999; Wang and Hill 1999; Wang *et al*. 1999). However, an accelerated rate of loss was observed when the level of dominance of deleterious mutations is sufficiently high and/or selection is sufficiently strong, due to the background selection (BGS) effect of deleterious mutations (Charlesworth *et al*. 1993), which causes a reduction in the effective population size experienced by linked neutral variants

AOD due to LD is an attractive potential explanation for the maintenance of unexpectedly high levels of variability in small populations. However, we lack a quantitative theory concerning the conditions under which it produces noticeable effects on levels of variability, other than for the cases of selfing and sib-mating lines studied by Wang and Hill (1999). It is also unclear when retardation versus acceleration of the rate of loss of variability is likely to prevail in randomly mating populations, especially with multiple loci subject to mutation and selection. As a first step toward such a theory, we present some analytical, numerical and simulation results on the simplest possible model: a neutral locus linked to another locus subject to selection. We also present simulation results for a neutral locus surrounded by a small number of selected loci. Selection can either involve partially recessive deleterious mutations or heterozygote advantage.

Our main focus is on a small, randomly mating population, founded from a large initial equilibrium population, mimicking experiments on the rates of loss of neutral variability in laboratory populations. We derive approximate expressions for the apparent selection coefficients against homozygotes at the neutral locus, as well for the rate of loss of neutral variability. We show that, contrary to what seems to have been widely assumed, these apparent selection coefficients have nothing to do with the rate of loss of variability. In particular, with selection against deleterious mutations there is a wide range of parameter space in which BGS accelerates the loss of variability, but an apparent selective advantage to heterozygotes at the neutral locus is always observed. When selection is very weak, AOD always causes a retardation of the loss of neutral variability, unless mutations are close to semidominance in their fitness effects.

## Theoretical Models and Methods

### Apparent selection coefficients against homozygotes at a neutral locus caused by a linked locus subject to mutation to deleterious alleles

We assume that the neutral locus is segregating for two alleles, A_1_ and A_2_, with frequencies *x* and *y* = 1 - *x*, respectively, in a given generation. The selected locus has wild-type and deleterious mutant alleles, B_1_ and B_2_, respectively, with frequencies *p* and *q = 1 - p*. The selection and dominance coefficients at this locus are *h* and *s*, such that the relative fitnesses of B_1_B_1_, B_1_B_2_, and B_2_B_2_ are 1, 1 - *hs* and 1 - *s*. The mutation rates for B_1_ to B_2_ and B_2_to B_1_ are *u* and *v*, respectively. Let the haplotype frequencies of A_1_B_1_, A_1_B_2_, A_2_B_1_ and A_2_B_2_ be *y*_1_, *y*_2_, *y*_3_, and *y*_4_. We have *x = y*_1_ + *y*_2_, *q* = *y*_2_ + *y*_4_. The frequency of recombination between the two loci is *c*.

With random mating, the apparent fitnesses (denoted by tildes) of the three genotypes A_1_A_1_, A_1_A_2_ and A_2_A_2_ at the neutral locus, conditional on these haplotype frequencies, are as follows (Sved 1968; Ohta 1971):

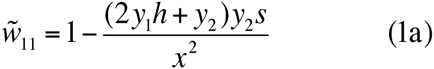

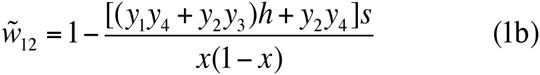

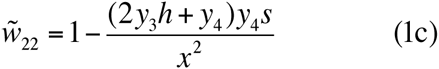

The extent of apparent selection at a neutral locus, induced by LD in a randomly mating finite population, can be assessed by determining the expectations of these apparent fitnesses over the distribution of haplotype frequencies induced by drift, conditioning on segregation at the A locus (Ohta and Kimura 1970; Ohta 1971). In the present case, the starting point is assumed to be a population of infinite size, at equilibrium under mutation and selection at the B locus and with no LD between the two loci. It is thereafter maintained at a population size of *N* breeding individuals each generation. For simplicity, a Wright-Fisher population model is assumed, so that *N* is also the effective population size.

The expectations for a given generation *t* after the foundation of the population are most simply found by expressing the haplotype frequencies in terms of the products of the relevant allele frequencies and the coefficient of linkage disequilibrium, *D = y*_1_ *y*_4_ - *y*_2_ *y*_3_. The expectation of *D*, E{*D*}, remains at zero, since selection, mutation and drift do not affect the direction of association between the two loci; this applies generally to *E{Dq*^*i*^}, where *i* is an arbitrary non-negative integer (a proof is given in section S4 of the Supplementary Information). This does not, however, imply that quantities involving the expectation of products of *D* and other functions of the allele frequencies at the two loci can be ignored, which complicates the analyses.

Simple algebra (Ohta 1971) shows that the expected apparent selection coefficients against A_1_A_1_ and A_2_A_2_ homozygotes (neglecting second-order terms in *s*) over a set of replicate populations that are segregating at the A locus are given by:

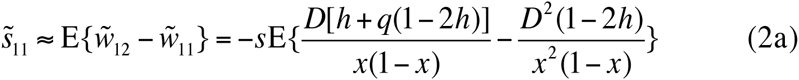

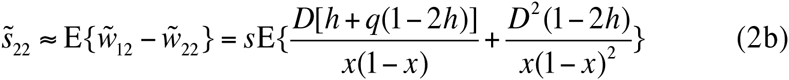

For simplicity, from now on we will simply refer to these as the apparent selection coefficients.

#### Approximations for the apparent selection coefficients

It has previously been assumed that the terms in *D* in these expressions can be neglected, so that the extent of apparent overdominance at the neutral locus is given by the terms in *D*_2_ alone. This allows approximations for the apparent selection coefficients to be derived; two different approaches have been used, as described by Sved (1968), Ohta and Kimura (1970) and Ohta (1971), and by Bierne *et al*. (2000), respectively. We will present an alternative approach here, which takes into account the initial allele frequency at the neutral locus in the founding generation, as well as the subsequent effects of drift in creating a probability distribution around this frequency.

The measures of apparent overdominance in Equations 2 can be conveniently be written as:

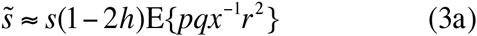

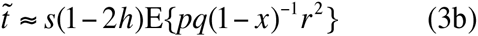

where *r*^2^ = *D*^2^/[*x(1-x)pq]* is the squared correlation coefficient in allelic state between the two loci (Hill and Robertson 1968). The difference between Equations 2 and 3 in the notation for the apparent selection coefficients is intended to emphasize the use of terms in *D* alone in Equations 3.

In order to obtain useful approximate expressions for these apparent selection coefficients at an arbitrary time *t*, we assume that the probability distributions of *x*, *q* and *r* are independent of each other, which is of course not exact. We can then write:

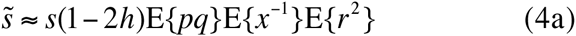

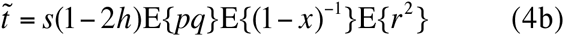

The quantity *s*(1 - *2h*)E{*pq*} is equal to the expectation of the inbred load, *B*, caused by the deleterious mutations, i.e., the expected difference in fitness between the logarithms of the mean fitness of a randomly mating population and of a completely homozygous population with the same allele frequencies (Greenberg and Crow 1960). Because of the loss of variability due to drift, E{*B*} will in general be smaller than the inbred load for an infinite population, *B* = p*q*s* (1 - *2h*), where *q\** is the equilibrium frequency of B_2_ under mutation and selection in an infinite population (Glémin *et al*. 2003). If *v* ≪ *u*, as assumed here, *q\** is given by the following expression (Crow and Kimura 1970, p.260):

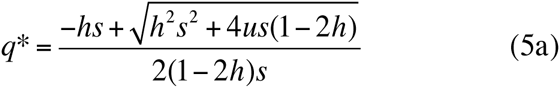

If *hs* is ≫ *u* and *h* > 0, this reduces to the familiar result of Haldane (1927):

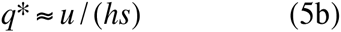

This yields the widely used formula for the inbred load in an infinite population:

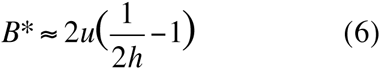

Weakly selected loci may contribute substantially to the inbred load and associative overdominance in the initial population, so that *q\** could be substantially greater than zero. The more general expressions for *q\** and *B\** will, therefore, be used here.

An exact analytic treatment of how E{*B*} changes over time in a finite population is, unfortunately, very difficult. Since the expected change in allele frequency due to selection is a third degree function of *q* when *h* ≠ 0.5, the third and higher moments of *q* enter into the expected change per generation in *B* under drift, mutation and selection, so that closed expressions cannot be obtained without using a full solution to the diffusion equation under selection with arbitrary *h*, (e.g., Balick *et al*. 2015). The simplest way to obtain tractable analytical results is to make the assumption that departures caused by drift from *q\** are sufficiently small that the change in allele frequency due to selection per generation, (∆_*s*_*q*), can be linearized around *q\**, and hence equated to (∆_*s*_*q*)_*q*\*_ + (*q - q*\*)(*d*∆_*s*_*q/dq*)_*q*\*_ (e.g., Charlesworth and Charlesworth 2010, p.355). If the mutational contributions to the change in allele frequency are included, the net change in *q* is equal to (*q - q*\*) (*d*∆_*s*_*q/dq*)_*q*\*_ - (*u + v*). The expected change in *q* in a finite population is then zero, and a recursion equation for the variance of *q, V*_*q*_, can easily be obtained, yielding a simple expression for the approximate value of E{*pq*} in a given generation (see Appendix).

An approximate linear recursion equation for the expectation of *r*^2^ can be obtained in a similar way, following the approach of Sved (1972) (see Appendix). The quantity σ^2^_*d*_ = E{D^2^}/E{*xypq*} is also frequently used as a descriptor of LD in finite populations, since it can be calculated by using diffusion equations (Ohta 1971; Ohta and Kimura 1971), as discussed below. While a simple recursion for σ^2^_*d*_ does not exist, a heuristic approach is to assume that it has a similar form to that for *r*^2^, replacing the equilibrium value of *r*_2_ given by Sved’s approach by the (smaller) equilibrium value of σ^2^_*d*_ (see Appendix). We can then substitute the expected value of σ^2^_*d*_ for the corresponding expectation of *r*_2_ in Equations 3. Both of these approaches ignore any effects of selection on LD.

We also need to obtain the expectations of *x*^−1^ and (1 - *x*)^−1^ for a given generation, conditioned on segregation at the neutral locus. This can be done using the diffusion equation solution for the probability distribution at a biallelic locus under pure drift (Kimura 1955), using the terms involving the first few eigenfunctions of the power series representation of the probability distribution, conditioned on an initial allele frequency *x*0 (see Appendix). The approximate expectations of *x*^−1^ and (1 - *x*)^−1^ can then easily be determined (see Appendix). As shown by Fisher (1930), this distribution is asymptotically close to a uniform distribution, with some slight deviations in the terminal classes, and with a mean allele frequency of 0.5 regardless of the value of *x*_0_. The asymptotic values of *x*^−1^ and (1- *x*)^−1^ can then be calculated, which both approach 5.18 asymptotically for the population size of 50 used in the numerical examples below (Equation A11); this implies that the apparent selection coefficients against each homozygote should approach the same asymptotic value, provided that any effects of mutation at the neutral locus can be ignored over the timescale under consideration.

### Apparent selection coefficients induced by linkage to a locus with heterozygote advantage

The same machinery can be used when the selected locus is segregating for a pair of alleles where the B_1_B_1_ and B_2_B_2_ homozygotes have fitnesses 1 - *s* and 1 - *t* relative to a fitness of 1 for B_1_B_2_:

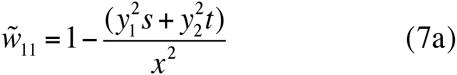

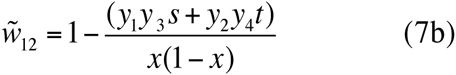

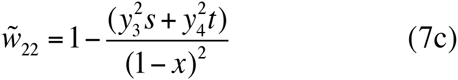

(Sved 1968; Ohta and Kimura 1970).

The approximate apparent selection coefficients are then given by:

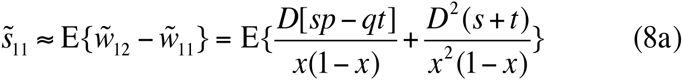

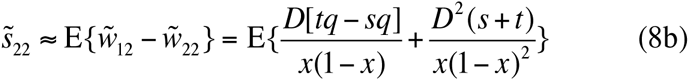

Using the same argument that led to Equations 4, these apparent selection coefficients can be approximated by:

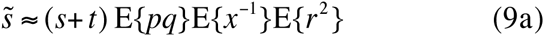

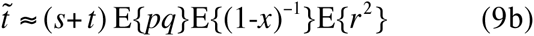

The expectation of the inbred load *B* in a finite population is now equal to (*s* + *t*) E{*pq*}, so that Equations 4 and 9 both involve the same function of E{*B*}. An approximation for E{*pq*} can be found by linearizing the recursion relation for *q* around the deterministic equilibrium, *q* = s/(s + t*), similarly to the procedure for the mutation-selection balance case. The coefficient *κ* that measures the rate of approach to equilibrium is now equal to *-st/(s + t*) (Charlesworth and Charlesworth 2010, p.355). In this case, no mutation is allowed at the locus under selection; this is because we are considering a single nucleotide site instead of a whole gene sequence, and mutations to new variants are unlikely to occur over the timescale with which we are concerned.

### The effect of selection at a linked locus on the amount of neutral variability

#### Lack of relevance of the apparent selection coefficients to the rate of loss of variability

It was assumed in previous theoretical studies that the apparent selection coefficients obtained by ignoring terms in *D* in Equations 2 and 8 provide a measure of the extent to which the loss of variability at a neutral locus, caused by drift, is retarded by linkage to a selected locus (Sved 1968; Ohta and Kimura 1970; Ohta 1971). But this assumption is mistaken, as can be seen as follows. Indeed, the apparent selection coefficients obtained from these equations may provide approximations for the apparent reductions in the fitnesses of the homozygotes at the neutral locus, but have no relation to the rate of loss of variability.

This can be seen as follows for the case of deleterious mutations at the B locus; a similar argument holds for the case of heterozygote advantage. The expected change in *x* given by Equations 2 (neglecting second-order terms in *s*) is:

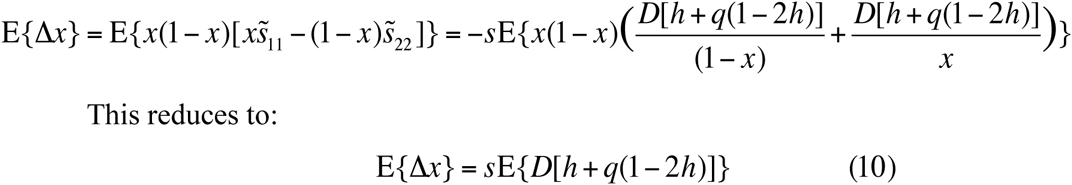

In other words, the terms in *D* contribute nothing to the change in the frequency of allele B_1_ at the neutral locus, which is determined entirely by the expected product of *D* and the additive effect of B_1_ on fitness, *a(q) = s[h + q(1 - 2h)]*. A result equivalent to this is used in models of selective sweeps on neutral allele frequencies at linked sites (Charlesworth and Charlesworth 2010, p.410). It is an example of the Price equation (Price 1970), since *Da* is the covariance between fitness and the allelic state with respect to A_1_ at the neutral locus (Santiago and Caballero 1995).

Since the change in the expectation of the heterozygosity, 2*x*(1 - *x*), at the neutral locus caused by the change in allele frequency due to selection at the B locus is approximated by the expectation of 2(1 - 2*x*)∆*x*, the accompanying change in the expected heterozygosity, *H* = 2E{*xy*}, is given by:

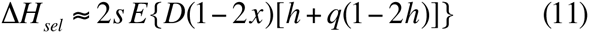

Similar results apply to the case of heterozygote advantage. Here, *a(q) = qt - ps = q(s+t) - s*, so that:

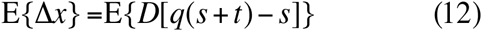

and

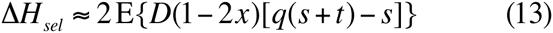

We cannot, therefore, use the apparent selection coefficients to assess the extent to which loss of variability at the neutral locus is affected by selection at a linked locus. Previous results using these selection coefficients (Ohta and Kimura 1970; Ohta 1971, 1973; Bierne *et al*. 2000) thus do not provide reliable measures of the extent to which AOD retards the loss of neutral variability.

#### How to calculate the rate of loss of variability

In order to solve this problem, we have extended the linear diffusion operator approach (Ohta and Kimura 1971; Ohta 1971) to include the effects of drift on allele frequencies at the selected locus, ignoring third-order and higher-order moments of *q* around *q\**. This requires the use of a 9-dimensional vector **Y** of functions of allele frequencies and *D*, with a corresponding 9 x 9 recursion matrix, **R** (see section 1 of the Supplementary Information for details). The elements of **Y** are as follows: *Y*_1_ = E{*xy*}, *Y*_2_ = E{*xyq*}, *Y*_3_ = E{*xyq*^2^}, *Y*_4_ = E{*D*(*x - y*)}, *Y*_5_ =E{*D(x - y)q*}, *Y*_6_ = E{*D(x - y)q*^2^}, *Y*_7_ = E{*D*^2^}, *Y*_8_ = E{*D*^2^*q*}, and *Y*_9_ = E{*D*^2^*q*^2^}.

Iteration of the **R** matrix provides an approximation for *σ*^2^_*d*_ in an arbitrary generation (which can used instead of E{*r*^2^} in Equations 3) as well as for *H*, since σ^2^_*d*_ = *Y*_7_/(*Y*_2_ - *Y*_3_) and *H = 2Y*_1_.

### Computer simulations

#### One neutral and one selected locus

The theoretical predictions described above were compared with Monte Carlo simulation results for a population started in mutation-selection balance at the selected locus and in linkage equilibrium with a neutral locus, and then run for a chosen number of generations at a fixed population size. In each generation, the new haplotype frequencies were calculated using the standard deterministic equations, and *2N* uniform random numbers were generated to sample each new haplotype from the cumulative distribution of haplotype frequencies.

Since the effects of a single selected locus on a linked neutral locus are very small, it was necessary to run a large number of replicate simulations (usually 10^7^) to obtain tight confidence intervals on the mean heterozygosity at the neutral locus

#### Multiple loci with selection and mutation

To simulate multiple loci subject to mutation and selection, we assumed additive fitness interactions across the relevant loci. For weak selection, this should yield very similar results to a multiplicative fitness model. The state of the population in a given generation is represented by the haplotype frequencies with respect to *n* loci subject to mutation and selection, following the same rules as in the single locus case, with a neutral locus in the centre of the chromosome. The recombination frequency between the loci under selection is *c*; the frequency of recombination between the neutral locus and each of the adjacent selected loci is *c*/2. Complete interference is assumed, so that only single crossovers are allowed, generating a frequency of recombination between the two terminal loci of (*n - 1)c*.

The frequencies of diploid genotypes at the start of a given generation are assumed to be given by random combinations the frequencies of the haplotypes determined in the previous generation, thus tacitly assuming an infinite number of new zygotes. The frequencies of haplotypes after recombination were calculated deterministically, using an extension of the three-locus algorithm described by Crow and Kimura (1970, p.51). The haplotype frequencies after mutation and selection were then calculated deterministically. (With weak effects of selection, mutation and recombination on genotype frequencies, the order in which these processes take place has a negligible effect on the final outcome.) The haplotype frequencies after drift were obtained by choosing *2N* haplotypes from a pool with the new haplotype frequencies, in the same way as above.

FORTRAN programs for all the cases mentioned here are available on request to BC.

## Results

### A single selected locus with mutation and selection

In order to generalize the results described below as far as possible, we have used the principle from diffusion equation theory that the outcome of the evolutionary process is determined by the products of the effective population size, *N*_*e*_, and the deterministic parameters, measuring time in units of 2*N*_*e*_ generations (Ewens 2004, p. 157). In the present case, where *N*_*e*_ = *N*, the simulation results for a given *N* and a set of mutation, selection and recombination parameters can be applied to a system with a population size of *KN* and deterministic parameters that are a factor of *K*^−1^ times those used here. For this reason, the selection and mutation parameters in most of the material described below are scaled by a factor of *2N*, thereby avoiding reference to a specific population size. We use *S = 2Ns, C = 2Nc, U= 2Nu* and *V = 2Nv* to denote these scaled parameters.

An additional point to note is that the assumption that the initial state of the population has *D* = 0 implies that only the first three elements of the initial **Y** vector are non-zero. Without loss of generality, the elements of **Y** can be divided by the product of the initial frequencies of A_1_ and A_2_, *x*_0_*y*_0_, and the resulting vector, **Y’**, can then be iterated using the **R** matrix. The ratio of the first element of **Y’** in generation *t* (1 - 1/2*N*)^t^ measures the ratio of the mean heterozygosity at the neutral locus relative to its value in the absence of selection at the B locus, which we denote by *H*_*rel*_. This procedure means that the matrix predictions for *H*_*rel*_ and *σ*^2^_*d*_ are independent of the initial allele frequencies at the neutral locus, since the initial value of **Y’** is independent of these frequencies. If, however, *D* were non-zero in the initial generation, only the asymptotic behaviour of the system would be independent of the initial allele frequencies.

#### Accuracy of the matrix approximation

We first consider the accuracy of the 9 x 9 matrix approximation. Figure 1 shows the values of *H*_*rel*_ and 2∆E{*xy*} = ∆*H*_*sel*_ from Equation 11 over *2N* generations, for two different dominance coefficients and with *C* = 0.1. We are assuming that selection and mutation at the B locus relates to deleterious mutations affecting the gene as a whole rather than individual nucleotide sites, whereas the alleles at the A locus represent a pair of variants chosen by an experimenter as neutral markers, and are not affected by mutation over the time course of the experiment. The selection and mutation parameters were chosen such that the equilibrium frequency of the deleterious mutation at the B locus was approximately 0.3. This may seem very high, but is consistent with the typical frequency per gene of putatively deleterious nonsynonymous mutations in *Drosophila* populations (Haddrill *et al*. 2010), although the selection coefficient and mutation rates used in the figure are probably much higher than is realistic.

**Figure 1.**
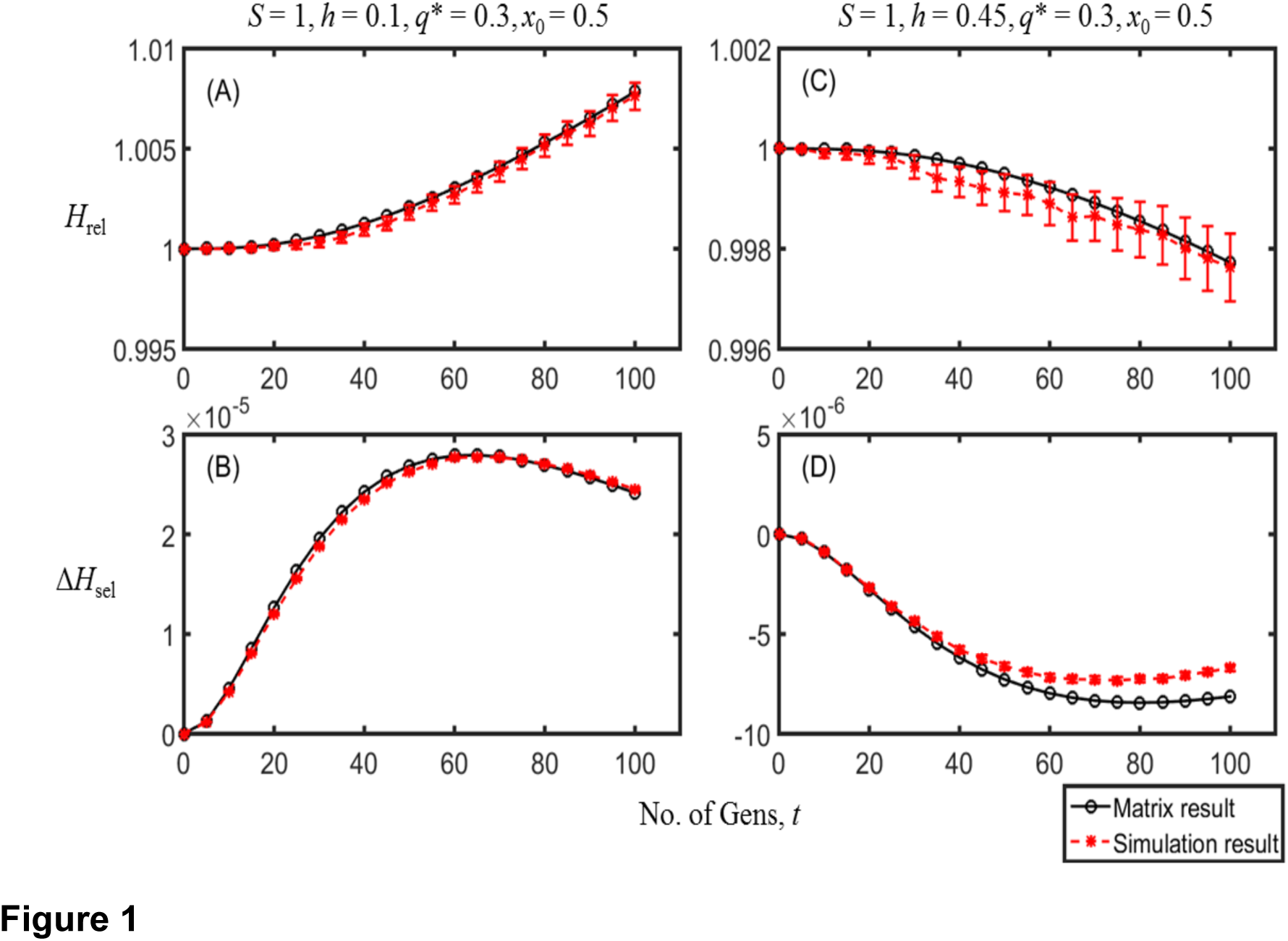
A comparison of the simulation results (red dots, with 95% confidence interval error bars) with the matrix approximation (black dots) for the case of mutation and selection. The scaled selection coefficient and recombination rates are *S* = 1 and *C* = 0.1. The population size is *N* = 50; the initial frequency of the neutral allele A_1_is *x*0 = 0.5; the dominance coefficient is *h* = 0.1 for A and B and *h* = 0.45 for Panels C and D. The mutation rates are such that the equilibrium frequency of the deleterious allele B_2_, *q\**, is approximately 0.3, with *v = 0*.1*u*. Panels A and C show the heterozygosity relative to neutral expectation at the A locus, *H*_*rel*_; Panels B and D show the change in heterozygosity per generation at the A locus due to selection at the B locus, ∆*H*_*sel*_. *h* = 0.1 and *h* = 0.45 are on opposite sides of the critical *h* value (approximately 0.37 for 2*N* generations), so that Panel A shows a retardation of loss of neutral variability, whereas Panel C shows an acceleration.

Agreement between the simulation and matrix results is remarkably good for this parameter set. A value of *h* = 0.1 is associated with an elevation of *H*_*rel*_ over one in later generations (retardation of loss of neutral variability), whereas *h* = 0.45 is associated with *H*_*rel*_ less than one (acceleration of loss of neutral variability). This illustrates the conflict between AOD and BGS, noted previously in simulation results (Pamilo and Palsson 1999; Wang and Hill 1999); for sufficiently low *h*, AOD is the dominant force as far as the level of neutral variability is concerned, whereas BGS dominates when *h* is sufficiently close to ½. We provide some insights into the reason for this behavior in the next but one section.

This conflict implies the existence of a critical value of *h, h*_*c*_, where the two effects exactly cancel each other; the properties of *h*_*c*_ are examined in more detail below. Unfortunately, there is no exact fixed value of *h*_*c*_, since the extent to which *H_rel_* is greater or less than one depends on the generation in question, as shown in Figure 2, which uses the matrix results. As *h* increases from 0 to 0.5, *H*_*rel*_ at both generation *N* and *2N* declines. The insert shows *h*_*c*_ is larger for the earlier generations. However, for the selection and mutation parameters used in Figure 2, *h*_*c*_ is close to 0.37 for all generations.

**Figure 2.**
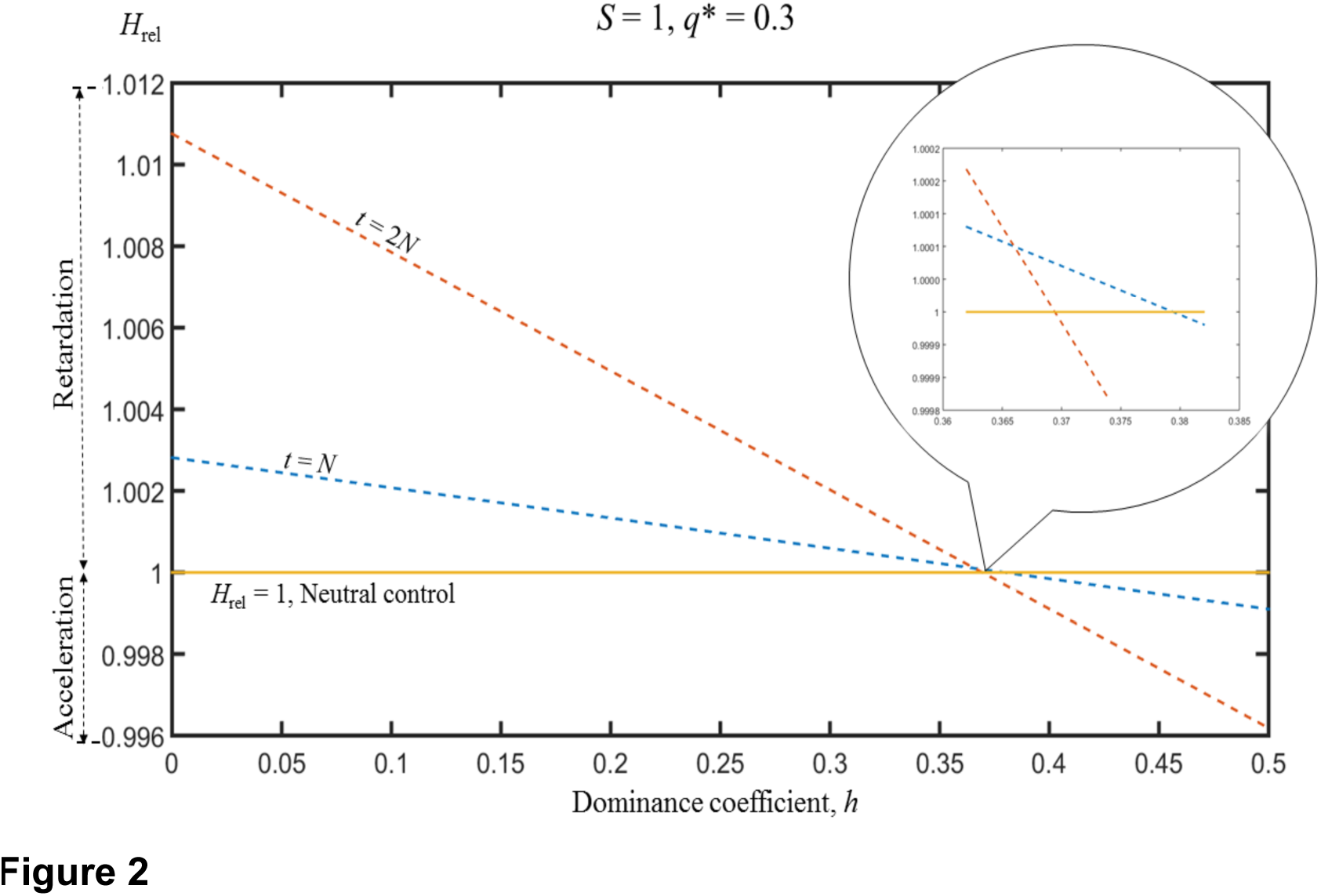
The heterozygosity at the A locus relative to neutral expectation, *H*_*rel*_, as a function of the dominance coefficient, *h*, at generations *N* and 2*N*, for the case of mutation and selection. *S* = 1, *C* = 0.1, *N* = 50 and *q\** = 0.3. The results were obtained using the matrix approximation.

When *q\** is small and *S* is of order 1 or less, agreement with the simulations is good only in earlier generations of the process, reflecting effects of such high a level of dispersion of allele frequencies at the B locus around *q\** that our neglect of higher-order terms is inaccurate. In general, however, the matrix gives reasonably good approximations up to *2N* generations, although the extent of retardation of the rate of loss of variability tends to be underestimated (or the rate of acceleration overestimated) by the matrix approximation with small *q\** (See Figure S2 in Supplementary Information, Section 9). We will often use results for this generation as a standard, since it is of similar order to the duration of the Drosophila experiments on AOD described in the Discussion.

#### Apparent selection coefficients and their relation to the rate of loss of variability

These conclusions are confirmed by the results displayed in Table 1, which also shows the simulation results for an additional measure of variability, the proportion of segregating neutral loci, *P*_*s*_, at 0.5*N* and *2N* generations, for a range of recombination rates. In addition, the exact and approximate apparent selection coefficients are displayed, together with the matrix and simulation values for *H*_*rel*_. It can be seen that the approximate apparent selection coefficients are similar in magnitude to the exact ones, but tend to exceed them, especially with very close linkage and for the neutral allele with the lower initial frequency. This is mainly because of the inaccuracy of the assumption of independence between the probability distributions of allele frequencies at the neutral and selected loci. Variability at the two loci is positively correlated, so that the expectation of *pqx*^−1^ in Equation 3a will tend to be smaller than E{*pq*}E{*x*^−1^} in Equation 4a.

**Table 1.**
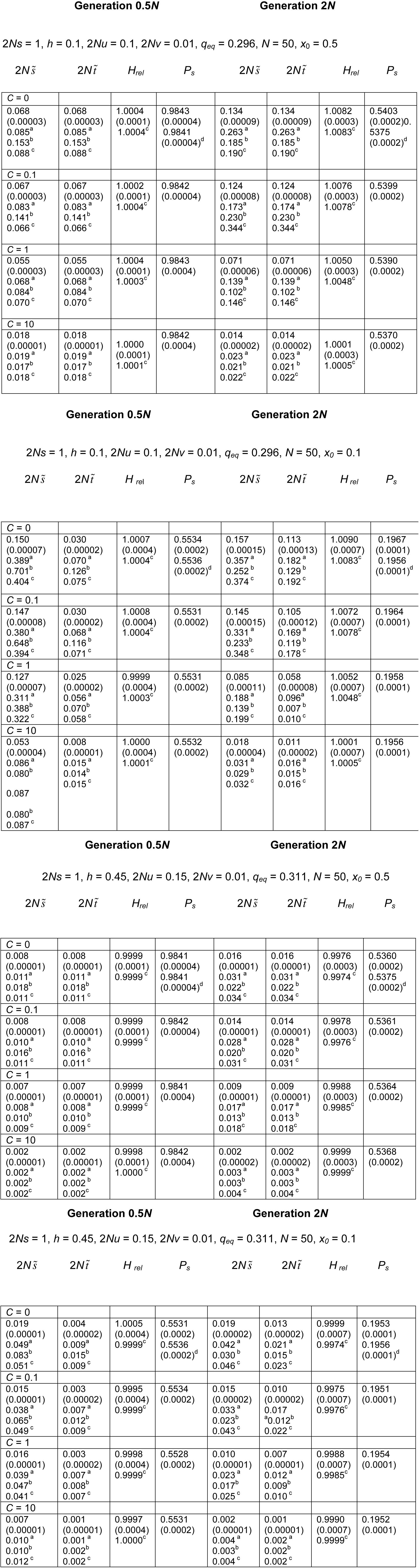
Simulation and theoretical results for one locus subject to mutation to deleterious alleles. ^a^ Approximation using neutral *r*_2_ recursion; ^b^ Approximation using neutral σ^2^*d* recursion; ^c^Approximation using matrix recursion with selection; ^d^ Expectation from neutral simulations. Standard errors for 10^7^ simulations are shown in parentheses.

| Generation $0.5N$ | | | | Generation $2N$ | | | |
| --- | --- | --- | --- | --- | --- | --- | --- |
| $2N\tilde{s}$ | $2N\tilde{t}$ | $H_{rel}$ | $P_s$ | $2N\tilde{s}$ | $2N\tilde{t}$ | $H_{rel}$ | $P_s$ |
| **C = 0** |  |  |  |  |  |  |  |
| 0.068<br>(0.00003)<br>0.085<sup>a</sup><br>0.153<sup>b</sup><br>0.088<sup>c</sup> | 0.068<br>(0.00003)<br>0.085<sup>a</sup><br>0.153<sup>b</sup><br>0.088<sup>c</sup> | 1.0004<br>(0.0001)<br>1.0004<sup>c</sup> | 0.9843<br>(0.00004)<br>0.9841<br>(0.00004)<sup>d</sup> | 0.134<br>(0.00009)<br>0.263<sup>a</sup><br>0.185<sup>b</sup><br>0.190<sup>c</sup> | 0.134<br>(0.00009)<br>0.263<sup>a</sup><br>0.185<sup>b</sup><br>0.190<sup>c</sup> | 1.0082<br>(0.0003)<br>1.0083<sup>c</sup> | 0.5403<br>(0.0002)0.5375<br>(0.0002)<sup>d</sup> |
| **C = 0.1** |  |  |  |  |  |  |  |
| 0.067<br>(0.00003)<br>0.083<sup>a</sup><br>0.141<sup>b</sup><br>0.066<sup>c</sup> | 0.067<br>(0.00003)<br>0.083<sup>a</sup><br>0.141<sup>b</sup><br>0.066<sup>c</sup> | 1.0002<br>(0.0001)<br>1.0004<sup>c</sup> | 0.9842<br>(0.00004) | 0.124<br>(0.00008)<br>0.173<sup>a</sup><br>0.230<sup>b</sup><br>0.344<sup>c</sup> | 0.124<br>(0.00008)<br>0.174<sup>a</sup><br>0.230<sup>b</sup><br>0.344<sup>c</sup> | 1.0076<br>(0.0003)<br>1.0078<sup>c</sup> | 0.5399<br>(0.0002) |
| **C = 1** |  |  |  |  |  |  |  |
| 0.055<br>(0.00003)<br>0.068<sup>a</sup><br>0.084<sup>b</sup><br>0.070<sup>c</sup> | 0.055<br>(0.00003)<br>0.068<sup>a</sup><br>0.084<sup>b</sup><br>0.070<sup>c</sup> | 1.0004<br>(0.0001)<br>1.0003<sup>c</sup> | 0.9843<br>(0.0004) | 0.071<br>(0.00006)<br>0.139<sup>a</sup><br>0.102<sup>b</sup><br>0.146<sup>c</sup> | 0.071<br>(0.00006)<br>0.139<sup>a</sup><br>0.102<sup>b</sup><br>0.146<sup>c</sup> | 1.0050<br>(0.0003)<br>1.0048<sup>c</sup> | 0.5390<br>(0.0002) |
| **C = 10** |  |  |  |  |  |  |  |
| 0.018<br>(0.00001)<br>0.019<sup>a</sup><br>0.017<sup>b</sup><br>0.018<sup>c</sup> | 0.018<br>(0.00001)<br>0.019<sup>a</sup><br>0.017<sup>b</sup><br>0.018<sup>c</sup> | 1.0000<br>(0.0001)<br>1.0001<sup>c</sup> | 0.9842<br>(0.0004) | 0.014<br>(0.00002)<br>0.023<sup>a</sup><br>0.021<sup>b</sup><br>0.022<sup>c</sup> | 0.014<br>(0.00002)<br>0.023<sup>a</sup><br>0.021<sup>b</sup><br>0.022<sup>c</sup> | 1.0001<br>(0.0003)<br>1.0005<sup>c</sup> | 0.5370<br>(0.0002) |
| **C = 0** |  |  |  |  |  |  |  |
| 0.150<br>(0.00007)<br>0.389<sup>a</sup><br>0.701<sup>b</sup><br>0.404<sup>c</sup> | 0.030<br>(0.00002)<br>0.070<sup>a</sup><br>0.126<sup>b</sup><br>0.075<sup>c</sup> | 1.0007<br>(0.0004)<br>1.0004<sup>c</sup> | 0.5534<br>(0.0002)<br>0.5536<br>(0.0002)<sup>d</sup> | 0.157<br>(0.00015)<br>0.357<sup>a</sup><br>0.252<sup>b</sup><br>0.374<sup>c</sup> | 0.113<br>(0.00013)<br>0.182<sup>a</sup><br>0.129<sup>b</sup><br>0.192<sup>c</sup> | 1.0090<br>(0.0007)<br>1.0083<sup>c</sup> | 0.1967<br>(0.0001)<br>0.1956<br>(0.0001)<sup>d</sup> |
| **C = 0.1** |  |  |  |  |  |  |  |
| 0.147<br>(0.00008)<br>0.380<sup>a</sup><br>0.648<sup>b</sup><br>0.394<sup>c</sup> | 0.030<br>(0.00002)<br>0.068<sup>a</sup><br>0.116<sup>b</sup><br>0.071<sup>c</sup> | 1.0008<br>(0.0004)<br>1.0004<sup>c</sup> | 0.5531<br>(0.0002) | 0.145<br>(0.00015)<br>0.331<sup>a</sup><br>0.233<sup>b</sup><br>0.348<sup>c</sup> | 0.105<br>(0.00012)<br>0.169<sup>a</sup><br>0.119<sup>b</sup><br>0.178<sup>c</sup> | 1.0072<br>(0.0007)<br>1.0078<sup>c</sup> | 0.1964<br>(0.0001) |
| **C = 1** |  |  |  |  |  |  |  |
| 0.127<br>(0.00007)<br>0.311<sup>a</sup><br>0.388<sup>b</sup><br>0.322<sup>c</sup> | 0.025<br>(0.00002)<br>0.056<sup>a</sup><br>0.070<sup>b</sup><br>0.058<sup>c</sup> | 0.9999<br>(0.0004)<br>1.0003<sup>c</sup> | 0.5531<br>(0.0002) | 0.085<br>(0.00011)<br>0.188<sup>a</sup><br>0.139<sup>b</sup><br>0.199<sup>c</sup> | 0.058<br>(0.00008)<br>0.096<sup>a</sup><br>0.007<sup>b</sup><br>0.010<sup>c</sup> | 1.0052<br>(0.0007)<br>1.0048<sup>c</sup> | 0.1958<br>(0.0001) |
| **C = 10** |  |  |  |  |  |  |  |
| 0.053<br>(0.00004)<br>0.086<sup>a</sup><br>0.080<sup>b</sup><br>0.087<br>0.080<sup>b</sup><br>0.087<sup>c</sup> | 0.008<br>(0.00001)<br>0.015<sup>a</sup><br>0.014<sup>b</sup><br>0.015<sup>c</sup> | 1.0000<br>(0.0004)<br>1.0001<sup>c</sup> | 0.5532<br>(0.0002) | 0.018<br>(0.00004)<br>0.031<sup>a</sup><br>0.029<sup>b</sup><br>0.032<sup>c</sup> | 0.011<br>(0.00002)<br>0.016<sup>a</sup><br>0.015<sup>b</sup><br>0.016<sup>c</sup> | 1.0001<br>(0.0007)<br>1.0005<sup>c</sup> | 0.1956<br>(0.0001) |
| **C = 0** |  |  |  |  |  |  |  |
| 0.008<br>(0.00001)<br>0.011<sup>a</sup><br>0.018<sup>b</sup><br>0.011<sup>c</sup> | 0.008<br>(0.00001)<br>0.011<sup>a</sup><br>0.018<sup>b</sup><br>0.011<sup>c</sup> | 0.9999<br>(0.0001)<br>0.9999<sup>c</sup> | 0.9841<br>(0.00004)<br>0.9841<br>(0.00004)<sup>d</sup> | 0.016<br>(0.00001)<br>0.031<sup>a</sup><br>0.022<sup>b</sup><br>0.034<sup>c</sup> | 0.016<br>(0.00001)<br>0.031<sup>a</sup><br>0.022<sup>b</sup><br>0.034<sup>c</sup> | 0.9976<br>(0.0003)<br>0.9974<sup>c</sup> | 0.5360<br>(0.0002)<br>0.5375<br>(0.0002)<sup>d</sup> |
| **C = 0.1** |  |  |  |  |  |  |  |
| 0.008<br>(0.00001)<br>0.010<sup>a</sup><br>0.016<sup>b</sup><br>0.011<sup>c</sup> | 0.008<br>(0.00001)<br>0.010<sup>a</sup><br>0.016<sup>b</sup><br>0.011<sup>c</sup> | 0.9999<br>(0.0001)<br>0.9999<sup>c</sup> | 0.9842<br>(0.00004) | 0.014<br>(0.00001)<br>0.028<sup>a</sup><br>0.020<sup>b</sup><br>0.031<sup>c</sup> | 0.014<br>(0.00001)<br>0.028<sup>a</sup><br>0.020<sup>b</sup><br>0.031<sup>c</sup> | 0.9978<br>(0.0003)<br>0.9976<sup>c</sup> | 0.5361<br>(0.0002) |
| **C = 1** |  |  |  |  |  |  |  |
| 0.007<br>(0.00001)<br>0.008<sup>a</sup><br>0.010<sup>b</sup><br>0.009<sup>c</sup> | 0.007<br>(0.00001)<br>0.008<sup>a</sup><br>0.010<sup>b</sup><br>0.009<sup>c</sup> | 0.9999<br>(0.0001)<br>0.9999<sup>c</sup> | 0.9841<br>(0.0004) | 0.009<br>(0.00001)<br>0.017<sup>a</sup><br>0.013<sup>b</sup><br>0.018<sup>c</sup> | 0.009<br>(0.00001)<br>0.017<sup>a</sup><br>0.013<sup>b</sup><br>0.018<sup>c</sup> | 0.9988<br>(0.0003)<br>0.9985<sup>c</sup> | 0.5364<br>(0.0002) |
| **C = 10** |  |  |  |  |  |  |  |
| 0.002<br>(0.00001)<br>0.002<sup>a</sup><br>0.002<sup>b</sup><br>0.002<sup>c</sup> | 0.002<br>(0.00001)<br>0.002<sup>a</sup><br>0.002<sup>b</sup><br>0.002<sup>c</sup> | 0.9998<br>(0.0001)<br>1.0000<sup>c</sup> | 0.9842<br>(0.0004) | 0.002<br>(0.00002)<br>0.003<sup>a</sup><br>0.003<sup>b</sup><br>0.004<sup>c</sup> | 0.002<br>(0.00002)<br>0.003<sup>a</sup><br>0.003<sup>b</sup><br>0.004<sup>c</sup> | 0.9999<br>(0.0003)<br>0.9999<sup>c</sup> | 0.5368<br>(0.0002) |
| **C = 0** |  |  |  |  |  |  |  |
| 0.019<br>(0.00001)<br>0.049<sup>a</sup><br>0.083<sup>b</sup><br>0.051<sup>c</sup> | 0.004<br>(0.00002)<br>0.009<sup>a</sup><br>0.015<sup>b</sup><br>0.009<sup>c</sup> | 1.0005<br>(0.0004)<br>0.9999<sup>c</sup> | 0.5531<br>(0.0002)<br>0.5536<br>(0.0002)<sup>d</sup> | 0.019<br>(0.00002)<br>0.042<sup>a</sup><br>0.030<sup>b</sup><br>0.046<sup>c</sup> | 0.013<br>(0.00002)<br>0.021<sup>a</sup><br>0.015<sup>b</sup><br>0.023<sup>c</sup> | 0.9999<br>(0.0007)<br>0.9974<sup>c</sup> | 0.1953<br>(0.0001)<br>0.1956<br>(0.0001)<sup>d</sup> |
| **C = 0.1** |  |  |  |  |  |  |  |
| 0.015<br>(0.00001)<br>0.038<sup>a</sup><br>0.065<sup>b</sup><br>0.049<sup>c</sup> | 0.003<br>(0.00002)<br>0.007<sup>a</sup><br>0.012<sup>b</sup><br>0.009<sup>c</sup> | 0.9995<br>(0.0004)<br>0.9999<sup>c</sup> | 0.5534<br>(0.0002) | 0.015<br>(0.00002)<br>0.033<sup>a</sup><br>0.023<sup>b</sup><br>0.043<sup>c</sup> | 0.010<br>(0.00002)<br>0.017<sup>a</sup><br>0.012<sup>b</sup><br>0.022<sup>c</sup> | 0.9975<br>(0.0007)<br>0.9976<sup>c</sup> | 0.1951<br>(0.0001) |
| **C = 1** |  |  |  |  |  |  |  |
| 0.016<br>(0.00001)<br>0.039<sup>a</sup><br>0.047<sup>b</sup><br>0.041<sup>c</sup> | 0.003<br>(0.00002)<br>0.007<sup>a</sup><br>0.008<sup>b</sup><br>0.007<sup>c</sup> | 0.9998<br>(0.0004)<br>0.9999<sup>c</sup> | 0.5528<br>(0.0002) | 0.010<br>(0.00001)<br>0.023<sup>a</sup><br>0.017<sup>b</sup><br>0.025<sup>c</sup> | 0.007<br>(0.00001)<br>0.012<sup>a</sup><br>0.009<sup>b</sup><br>0.010<sup>c</sup> | 0.9988<br>(0.0007)<br>0.9985<sup>c</sup> | 0.1954<br>(0.0001) |
| **C = 10** |  |  |  |  |  |  |  |
| 0.007<br>(0.00001)<br>0.010<sup>a</sup><br>0.010<sup>b</sup><br>0.012<sup>c</sup> | 0.001<br>(0.00001)<br>0.001<sup>a</sup><br>0.002<sup>b</sup><br>0.002<sup>c</sup> | 0.9997<br>(0.0004)<br>1.0000<sup>c</sup> | 0.5531<br>(0.0002) | 0.002<br>(0.00001)<br>0.004<sup>a</sup><br>0.003<sup>b</sup><br>0.004<sup>c</sup> | 0.001<br>(0.00001)<br>0.002<sup>a</sup><br>0.002<sup>b</sup><br>0.002<sup>c</sup> | 0.9990<br>(0.0007)<br>0.9999<sup>c</sup> | 0.1952<br>(0.0001) |

With an initial frequency of A_1_ of 0.5, the apparent selection coefficients for A_1_A_1_ and A_2_A_2_ are always equal, as expected from Equations 3 and 4. In contrast, for an initial frequency of 0.1 there is initially a stronger apparent selection coefficient against A_1_A_1_, again as expected, but the selection coefficients converge on equality as the mean neutral allele frequency at segregating sites approaches 0.5; these are also similar to the selection coefficients for the case of an initial frequency of 0.5. The apparent selection coefficients at generation *2N* are quite close to the asymptotic values given by Equations 4 when the B locus has reached mutation-selection-drift equilibrium, if E{*pq*} is calculated using the linear approximation described above. For example, with *h* = 0.1, *C* = 0.1 and the selection and mutation parameters in Table 1, the asymptotic apparent selection coefficient for populations segregating at the A locus is 0.0014 when the equilibrium neutral value of σ_*d*_^2^ is used as the estimate of E{*r*_2_} in Equations 4, whereas the observed value at 2*N* generations for the case with *x*_0_= 0.5 is 0.0013. In contrast, the asymptotic value predicted by Equation 7 of Ohta (1971) is 0.0024, a somewhat worse fit. This probably reflects the fact that Ohta assumed a fixed allele frequency at the neutral locus, as well as *q* ≪* 1.

As expected, the magnitudes of the apparent selection coefficients decline sharply as the recombination rate increases, but always indicate heterozygote advantage, even with *C* = 10 and *h* = 0.45. But with *h* = 0.45, there is an acceleration of the loss of variability at the neutral locus when linkage is tight, despite the apparent selective advantage to heterozygotes. This confirms the above conclusion that the apparent selection coefficients have no relation to the extent to which mutation and selection at one locus affect the rate of loss of variability at a linked neutral locus, and show heterozygote advantage even when BGS is causing an accelerated loss of variability. Furthermore, if *C* is ≫ 1, little effect of selection on the rate of loss of variability can be detected (Table 1).

#### Conflict between AOD and BGS with respect to the rate of loss of variability

The existence of a conflict between AOD and BGS, and its evident dependence on the dominance coefficient, raises the question of how to make generalizations about the regions of parameter space in which one or other force is dominant. One way of approaching this problem is to study the properties of the leading eigenvalue of the **R** matrix, *λ*_0_, as a function of the relevant variables. If we equate *λ*_0_ to 1 - 1/(2*N*_*e*_), where *N*_*e*_ is the effective population size in the presence of selection at the B locus, 1/(1 - *λ*_0_) provides a measure of 2*N*_*e*_. Consider first the limiting case when *S* = 2*Ns* ≫ 1 and *u* ≪ *hs*, so that the expected frequency of A_2_ is close to *u/(hs*) (Equation 5b). The approximation for *λ*_0_ - 1 for this case (Equations S17 and S18 of section 2 of the Supplementary Information) shows that *N*_*e*_ is given by the standard equation for BGS at a single locus (Charlesworth and Charlesworth 2010, p.402), which relies on these conditions.

As argued by Whitlock and Barton (1997), the value of 2*N*_*e*_ obtained from 1/(*λ*_0_ - 1) gives the asymptotic value of the mean coalescent time, *T*_2_, for a pair of alleles at the neutral locus. Under the infinite sites model, the nucleotide diversity at mutation-drift equilibrium is equal to the product of the neutral mutation rate and 2*N*_*e*_ (Charlesworth and Charlesworth 2010, p.211), so that we can use these results to predict the effect of selection at the B locus on the equilibrium level of variability at the neutral locus. (See section S2 of the Supplementary Information, Equation S16, for a more rigorous derivation of this result in the present context.)

However, since the matrix approximation is inaccurate for later generations with *S* < 1 and small *q\**, this use of the leading eigenvalue to estimate *N*_*e*_ is limited in scope, and is not necessarily accurate for general *q\** and small *S*. The approximate approach described in the next section deals with the case when *S* is of order 1 or less.

#### Approximation for the rate of loss of variability with weak selection

The starting point for this analysis is Equation 11, which shows that the change in expected heterozygosity due to selection, ∆*H*_*sel*_, is the sum of two terms, ∆_1_ = −2*shY*_4_ and ∆_2_ = −2*s*(1 - 2*h*)*Y*_5_, where *Y*_4_ = E{*D*(*x - y*)} and *Y*_5_ = E{*D(x - y)q*}. We know from previous work on BGS that an acceleration of loss of variability always occurs in haploid models with deleterious mutations and drift; these are equivalent to diploid models with *h* = 0.5, in which case ∆_2_ is necessarily zero, and the process is entirely driven by ∆_1_; a necessary and sufficient condition for acceleration in this case is thus ∆_1_ < 0. This suggests that with *h* < 0.5 there may be a conflict between the two terms, whenever ∆_1_ < 0 and ∆_2_ > 0. Larger *h* gives more weight to ∆_1_ relative to ∆_2_, so that selection is more likely to reduce variability when *h* is large, in agreement with the numerical results just described. It is, however, unclear from this argument why small *S* should favour AOD over BGS.

The complexity of the recursion relations means that it is impossible to obtain simple, exact expressions for *Y*_4_ and *Y*_5_, but useful approximate results for the case of weak selection (*S* < 1) can be obtained as follows. Using expressions S4d and S4e in the Supplementary Information, the changes per generation in these moments in the absence of selection are:

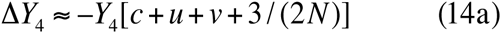

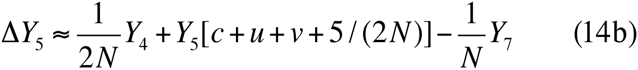

In the absence of selection, *Y*_4_ thus tends to zero, and *Y*_5_ tends to - *Y*_7_/[*N(C + U + v*) + 5], which is < 0 and < *Y*_7_ in magnitude. This suggests that the magnitude of ∆_1_ will be much smaller than that of ∆_2_ when selection is weak. This is confirmed by the following argument. When there is selection, approximate recursions for *Y*_4_ and *Y*_5_ can be obtained using the matrix approach described in section 1 of the Supplementary Information (Equations S5). We make the further simplification of dropping terms involving *u, v* and products of *s* or *D*_2_ with *q* and *q*_2_, on the assumption that these are small relative to similar terms involving *s* or *D*^2^. These assumptions will be violated when selection is strong in relation to drift (*S* > 1) or when *q\** is ≫ 0. In addition, for the neutral case, it is possible to show that *Y*_6_ and *Y*_5_ approach equality and *Y*_8_ approaches *Y*_7_/2 if *q\** is set to 0.5 (we neglect *Y*_9_, since it is considerably smaller than *Y*_8_): see Equations S31 of the Supplementary Information, section 6. Our numerical results show that these relations are approximately true under much wider conditions, so that we can replace *Y*_6_ with *Y*_5_, and *Y*_8_ with *Y*_7_/2 in Equations S5, allowing us to obtain a coupled pair of difference equations for *Y*_4_ and *Y*_5_ for a given value of *Y*_7_.

It is also convenient to rescale all terms by multiplying by *2N*, implying a new time-scale of *2N* generations. We can then write:

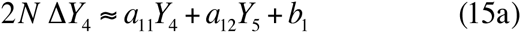

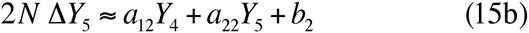

where

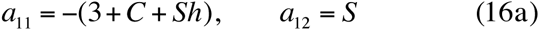

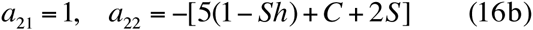

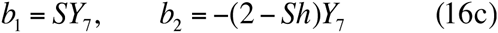

If the changes in *Y*_4_ etc. are sufficiently slow, we can set ∆*Y*_4_ and ∆*Y*_5_ to zero, and solve the resulting linear equations to obtain approximate expressions for *Y*_4_ and *Y*_5_ as functions of *Y*_7_:

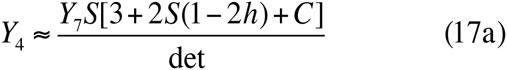

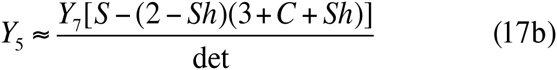

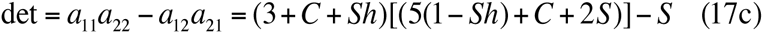

This shows that *Y*_4_ is *O*(*S*), whereas *Y*_5_ is of order one, so that ∆*H*_*sel*_ is dominated by ∆_1_ when *S* is small, provided that *h* < ½. The determinant is always positive for *S ≤* 60; this condition is not very restrictive given our assumption of relatively small *S*, and we will assume positivity from now on. Given that *X*_7_ > 0, it can easily be shown that *Y*_4_ > 0 when *h* < ½; *S* < 2 is sufficient for *Y*_5_ < 0 when *h* = 0.5. For small values of *h* these conditions are less stringent; with *h* = 0, *Y*_4_ > 0 independently of *S*, and *Y*_5_ < 0 when *S* < 6. Furthermore, neither *Y*_4_ nor *Y*_5_ depend on *q\**, except through their common factor of *Y*_7_. Our numerical results show that these conclusion are correct, provided that *S* is of order 1.

From Equation 11, the sign of the change due to selection in the heterozygosity at the neutral locus, ∆*H*_*sel*_, is the same as the sign of -[*h Y*_4_ + (1 - *2h*) *Y*_5_]. Variability will be increased by selection if ∆*H*_*sel*_ > 0 and reduced if ∆*H*_*sel*_ < 0, so that these results imply that the critical upper value of *h, h*_*c*_, required for an increase in neutral variability due to AOD, is also independent of *q\** under the conditions we have assumed. The value of *h*_*c*_ can be found by setting ∆*H*_*sel*_ = 0. Using Equations 11 together with the numerators of Equations 17a and 17b, we find that *h*_*c*_ is the value of *h* that satisfies the following cubic equation in *h*:

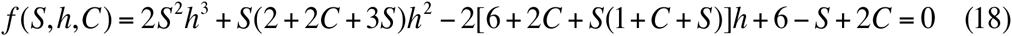

With *S* close to zero, Equation 18 reduces to 12*h* - 6 = 0, i.e. *h*_*c*_ = ½, suggesting that variability will always be increased for realistic *h* values when selection is very weak. For other cases, some insights into the properties of *h*_*c*_ can be obtained by assuming *C* = 0, which approximates cases with low recombination rates. With *S* = 1, the solution to the cubic is approximately *h*_*c*_ = 0.36, slightly smaller than the value of 0.37 obtained from the numerical results. Figure 3 shows a comparison of the values obtained from Equation 18 using Matlab symbolic calculations with the values from the matrix approximation for generation 2*N* with *C* = 0.1 (very similar patterns are obtained for *C* values of up to 1). It will be seen that they agree quite well when *S* < 1, regardless of the value of *q\**, but Equation 18 mostly underestimates *h*_*c*_ as *S* increases beyond 1, except for values of *q\** much smaller or much greater than 0.1. For *S* ≪ 1, dominance coefficients close to ***V2*** are compatible with an enhanced level of neutral variability, but BGS always prevails for values of *S* ≫ 3, even when *h* is very close to zero.

**Figure 3.**
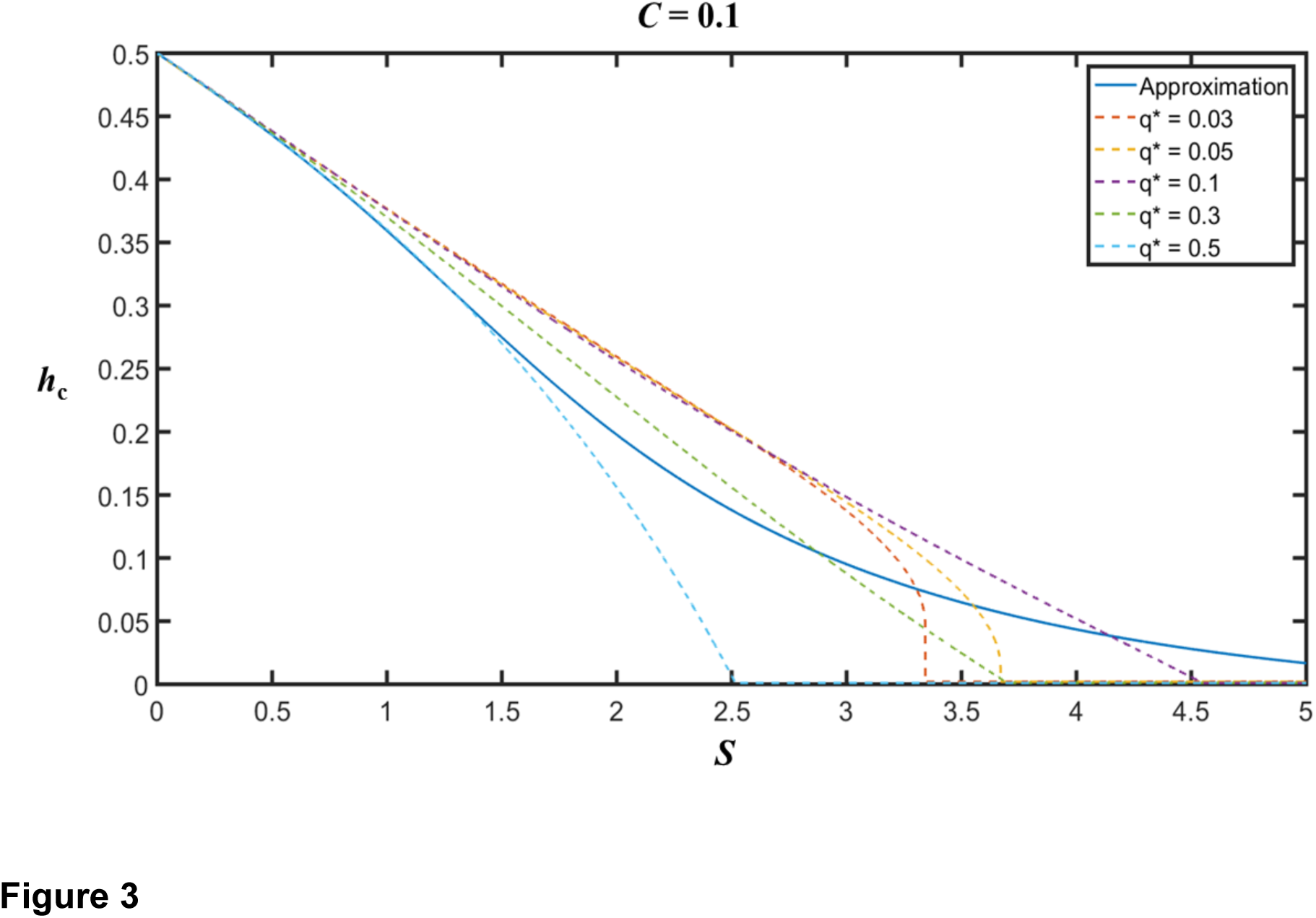
The critical dominance coefficient, *h*_*c*_, as a function of the scaled selection coefficient *S*, for the case of mutation and selection with *C* = 0.1 and *N* = 50. The initial frequency of A_1_was 0.5. The blue line is the value of *h*_*c*_ given by Equation 18. The dashed lines represent the values obtained from the matrix approximation at generation 2*N*, for several different values of the equilibrium frequency, *q\**, of the deleterious allele at the B locus.

The quantity *2N* ∆*H*_sel_/*H* is equivalent to *ε = N*_*e*_/*N* - 1, where *N*_*e*_ is the effective population size for the generation in question. For small *S*, we can approximate ε by a Taylor’s series around *S* = 0. We have:

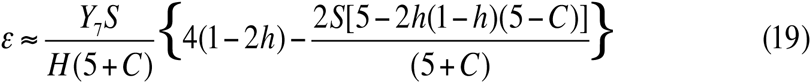

The first term in this equation is derived entirely from the contribution of *Y*_5_ to ∆*H*_*sel*_, since the contribution from *Y*_4_ is *O*(*S*_2_). This implies that selection will cause increased variability if *h* is slightly less than ½ when higher order terms in *S* are negligible, consistent with the conclusions reached above. With very weak selection, BGS thus does not operate unless mutations are close to being semidominant. The second-order term is always < 0 and increases in magnitude as *S* increases; under the most favourable situation (*h* = 0), *S* < 4(5 + *C*)/10 is necessary for *ε* > 0, which is slightly more restrictive than is indicated by the results in Figure 3. Figure 4 shows some examples of the extent to which this approximation match the simulation results; in general, it tends to underestimate the extent to which loss of heterozygosity is retarded by selection when *S*.

**Figure 4.**
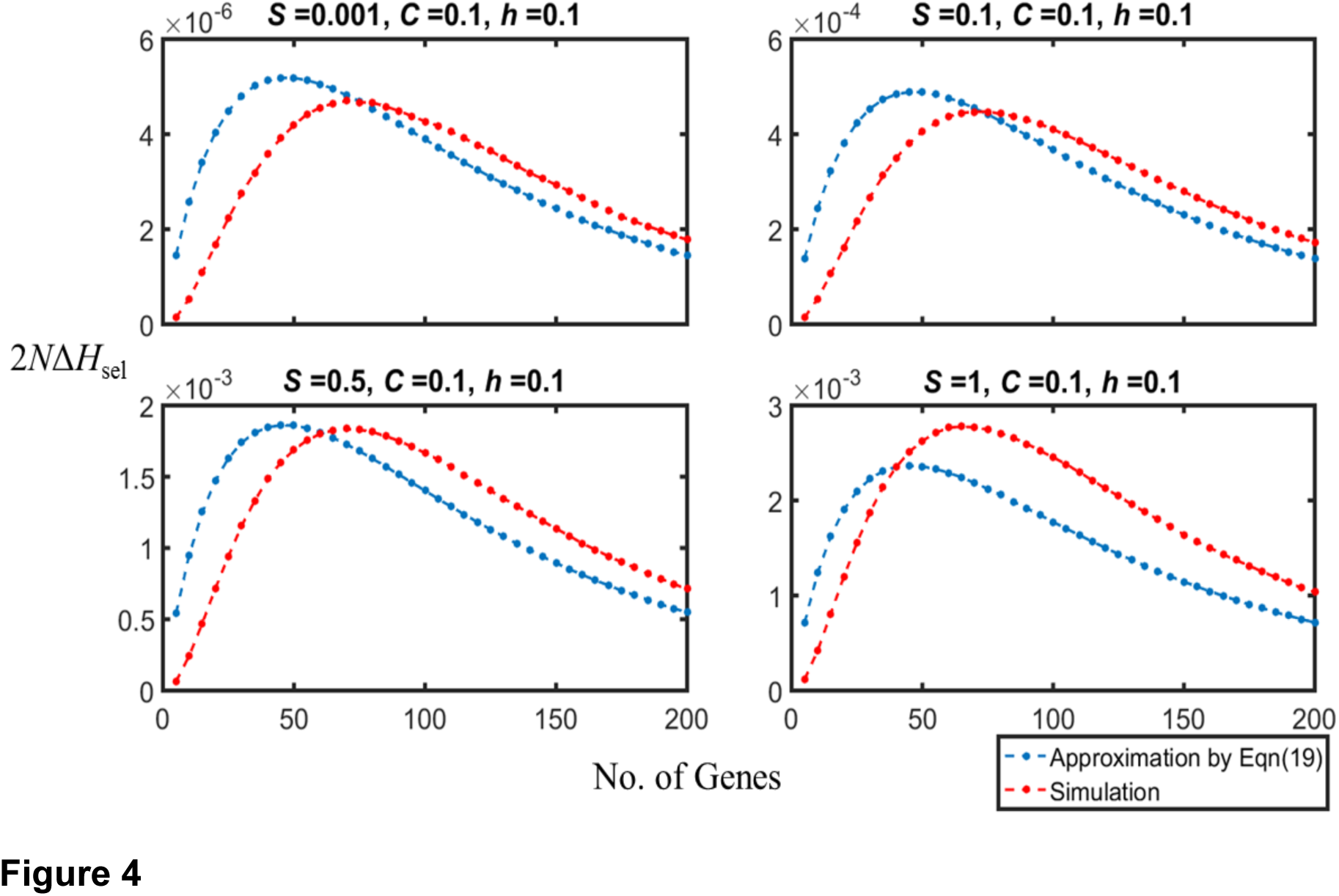
A comparison of the values of 2*N* ∆*H*_*sel*_ obtained from the simulations and from Equation 19 (using values of *Y*7 = E{D^2^} obtained from the simulations), with *C* = 0.1, *q\** =0.3 and *N* = 50. The initial frequency of A_1_was 0.5. The panels display the approximate results from Equation 19 (blue dots) and the simulation results (red dots) or different values of *S*. The confidence intervals are too small to be easily seen.

These approximations also imply that the magnitude of ∆*H*_*sel*_ is proportional to *Y*_7_ = E{*D*^2^}. If we approximate E{*D*^2^} by E{*xy*}E{*pq*}σ^2^_*d*_ = *H E{pq} σ*^2^_*d*_/*2*, as was done when obtaining the Equations 4 for the apparent selection coefficients, Equations 17 and 18 give:

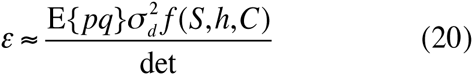

Since σ^2^_*d*_ E{*pq*}is ≤ 0.25, the quadratic dependence of the denominator on *C*, and the inverse relation between *σ*^2^_*d*_ and *C*, implies that there will only be a very small effect of selection on neutral variability when *C* ≫ 1, unless *S* ≫ *C*. In addition, the size of the effect is strongly determined by *S* and by E{*pq*}, since *σ*^2^_*d*_ takes a value between 0 and 1.

Overall, these results suggest that retardation of loss of neutral variability caused by a single selected locus is unlikely to be important except for *S* values of the order of 1. In such cases, if linkage is sufficiently tight, there can be a significant retardation of loss of neutral variability, and an enhancement of the equilibrium level of variability even when *h* approaches ½.

### Two selected loci with mutation and selection

From simulation studies of the effect of multiple, linked selected loci on the behaviour of neutral loci located among them, we would expect to see larger values of both the apparent selection coefficients and the level of neutral heterozygosity than with a single selected locus (Ohta 1971; Latter 1998; Pamilo and Palsson 1998, 1999; Wang and Hill 1999; Bierne *et al*. 2000). However, it is unclear whether the effects of multiple selected loci are approximately additive, or whether there is some degree of synergism.

With small population size, very close linkage, strong selection and a sufficiently high level of recessivity of deleterious mutations, simulations have shown that ‘crystallization’ of the population into two predominant and complementary haplotypes (+ − + − … and − + − +…, where + denotes wild-type and − mutant) with respect to the selected loci can occur, such that its behavior is very similar to that of a single locus with strong heterozygote advantage (Charlesworth and Charlesworth 1997; Pamilo and Palsson 1999; Palsson 2001). This can result in long-term maintenance of variability at the selected loci. This is strongly suggestive of a synergistic effect of multiple selected loci.

No analytical treatment of more than one selected locus has been done, so we have investigated this problem using simulations of a small number of selected loci. The results of simulations with two selected loci surrounding a neutral locus are shown in Table 2, using the same set of recombination frequencies between the neutral and nearest selected locus as in Table 1. As far as the apparent selection coefficients are concerned, with *h* = 0.1 and the lower two recombination rates, there is clear evidence that the apparent selection coefficients in generation 2*N* (but not necessarily in generation 0.5*N*) are somewhat larger than twice the apparent selection coefficients in the single locus case, indicating a degree of synergism. A similar pattern is found with four selected loci (results not shown).

**Table 2.**
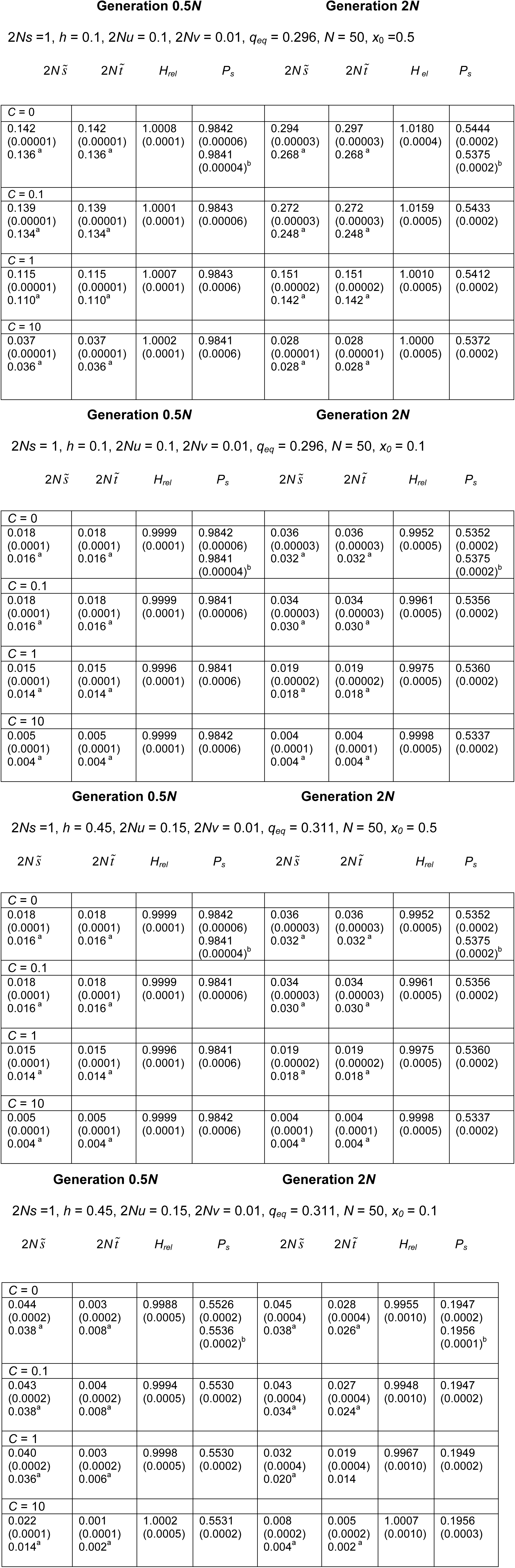
Simulation results for two loci subject to mutation to deleterious alleles. ^a^ Single locus simulation values multiplied by 2; ^b^ Expectation from neutral simulations. Standard errors for 5 x 10^6^ simulations with 2 selected loci are shown in parentheses.

It is more difficult to quantify the effect on *H*_*rel*_, since this is strongly dependent on the generation in question. With close linkage and *h* = 0.1, the values of *H*_*rel*_ - 1 in generation 2*N* in Table 2 are slightly larger than twice the corresponding values in Table 2. Conversely, with *h* = 0.45 and close linkage, for which *H*_*rel*_ < 1, the absolute values of *H*_*rel*_ - 1 in generation 2*N* are less than twice the corresponding values in Table 1, although the apparent selection coefficients are slightly larger than twice the values with a single selected locus. A more rigorous test for additivity is described in section S5 of the Supplementary Information; it suggests that the effects of two loci on *N_e_/N* - 1, as estimated from ∆*H*_*sel*_, are somewhat greater than additive when there is a retardation of loss of variability, and less than additive when there is an acceleration.

### A single selected locus with heterozygote advantage

The methods used for the case of mutation and selection can be used to model the case of linkage to a single selected locus with heterozygote advantage, simply by making appropriate modifications to the terms in the **R** matrix, and to the expressions for the changes in haplotype frequencies in the computer simulations (see Equations 9 and section 3 of the Supplementary Information). Figure 5 and Table 3 compare some examples of matrix and simulation results; these again suggest that the matrix predictions are accurate for this parameter range. Once again, the ratio of recombination to the strength of selection has to be sufficiently small for a substantial effect.

**Figure 5.**
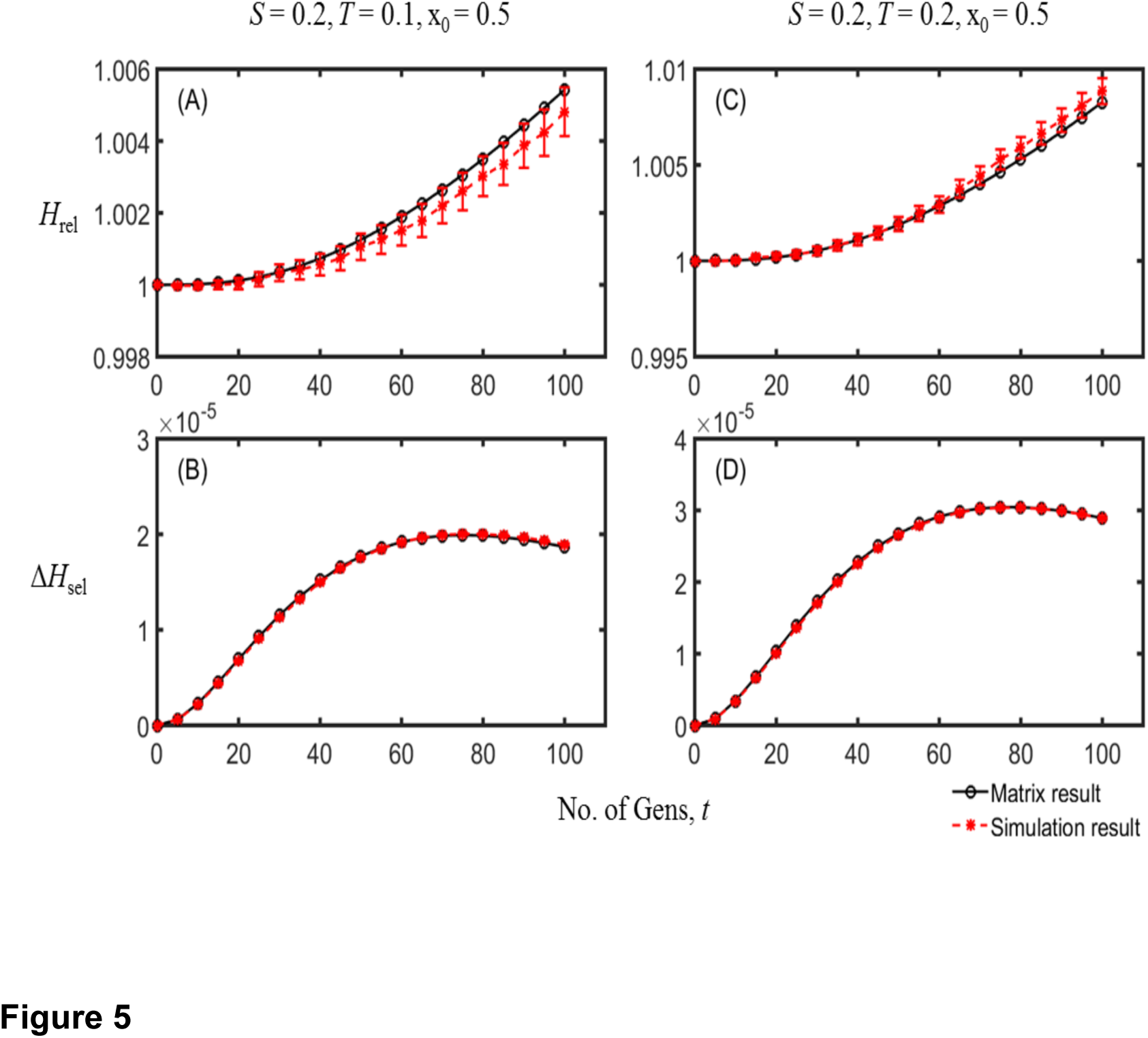
A comparison of the simulation results (red dots, with 95% confidence interval error bars) with the matrix approximation (black dots) for the case of heterozygote advantage. The left-hand panels plot *H*_*rel*_ and ∆*H*_*sel*_ against time for scaled selection coefficients *S* = 0.2 and *T* = 0.1; the right-hand panels give the values of these variables when *S* = 0.2 and *T* = 0.2. *N* = 50, *C* = 0.1, and the initial frequency of A_1_was 0.5.

**Table 3.**
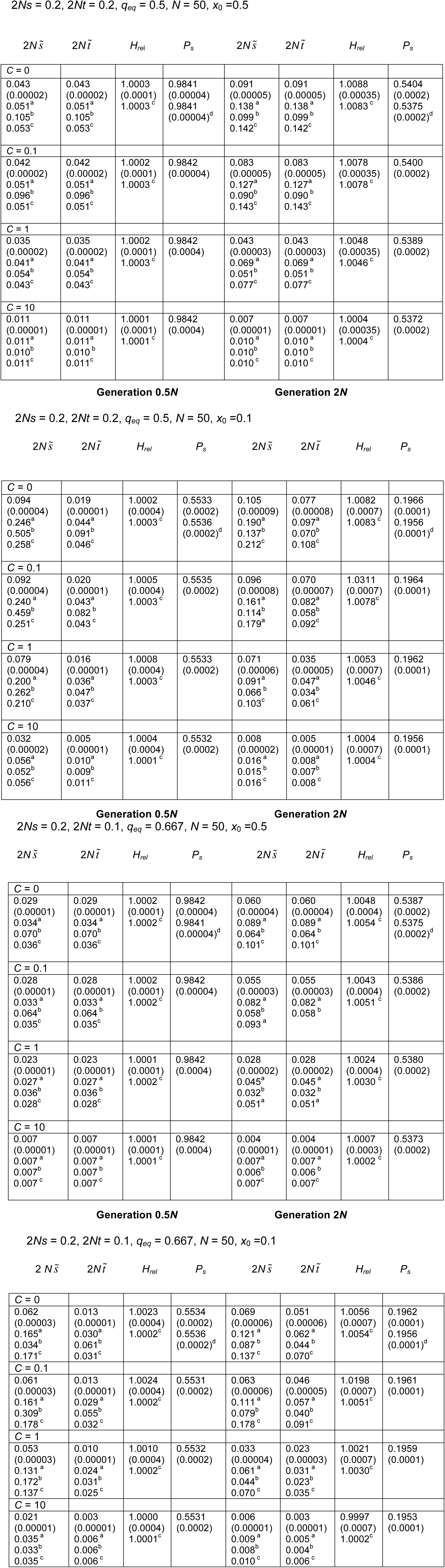
Simulation and theoretical results for a single locus with heterozygote advantage. ^a^ Approximation using neutral *r*_2_ recursion; ^b^ Approximation using neutral *σ^2^d* recursion; Approximation using matrix recursion with selection; Expectation from neutral simulations. Standard errors for 10^7^ simulations are shown in parentheses.

| $2N\tilde{s}$ | $2N\tilde{t}$ | $H_{rel}$ | $P_s$ | $2N\tilde{s}$ | $2N\tilde{t}$ | $H_{rel}$ | $P_s$ |
| --- | --- | --- | --- | --- | --- | --- | --- |
| **C = 0** |  |  |  |  |  |  |  |
| 0.043<br>(0.00002) | 0.043<br>(0.00002) | 1.0003<br>(0.0001) | 0.9841<br>(0.00004) | 0.091<br>(0.00005) | 0.091<br>(0.00005) | 1.0088<br>(0.00035) | 0.5404<br>(0.0002) |
| 0.051<sup>a</sup> | 0.051<sup>a</sup> | 1.0003<sup>c</sup> | 0.9841<br>(0.00004)<sup>d</sup> | 0.138<sup>a</sup> | 0.138<sup>a</sup> | 1.0083<sup>c</sup> | 0.5375<br>(0.0002)<sup>d</sup> |
| 0.105<sup>b</sup> | 0.105<sup>b</sup> |  |  | 0.099<sup>b</sup> | 0.099<sup>b</sup> |  |  |
| 0.053<sup>c</sup> | 0.053<sup>c</sup> |  |  | 0.142<sup>c</sup> | 0.142<sup>c</sup> |  |  |
| **C = 0.1** |  |  |  |  |  |  |  |
| 0.042<br>(0.00002) | 0.042<br>(0.00002) | 1.0002<br>(0.0001) | 0.9842<br>(0.00004) | 0.083<br>(0.00005) | 0.083<br>(0.00005) | 1.0078<br>(0.00035) | 0.5400<br>(0.0002) |
| 0.051<sup>a</sup> | 0.051<sup>a</sup> | 1.0003<sup>c</sup> |  | 0.127<sup>a</sup> | 0.127<sup>a</sup> | 1.0078<sup>c</sup> |  |
| 0.096<sup>b</sup> | 0.096<sup>b</sup> |  |  | 0.090<sup>b</sup> | 0.090<sup>b</sup> |  |  |
| 0.051<sup>c</sup> | 0.051<sup>c</sup> |  |  | 0.143<sup>c</sup> | 0.143<sup>c</sup> |  |  |
| **C = 1** |  |  |  |  |  |  |  |
| 0.035<br>(0.00002) | 0.035<br>(0.00002) | 1.0002<br>(0.0001) | 0.9842<br>(0.0004) | 0.043<br>(0.00003) | 0.043<br>(0.00003) | 1.0048<br>(0.00035) | 0.5389<br>(0.0002) |
| 0.041<sup>a</sup> | 0.041<sup>a</sup> | 1.0003<sup>c</sup> |  | 0.069<sup>a</sup> | 0.069<sup>a</sup> | 1.0046<sup>c</sup> |  |
| 0.054<sup>b</sup> | 0.054<sup>b</sup> |  |  | 0.051<sup>b</sup> | 0.051<sup>b</sup> |  |  |
| 0.043<sup>c</sup> | 0.043<sup>c</sup> |  |  | 0.077<sup>c</sup> | 0.077<sup>c</sup> |  |  |
| **C = 10** |  |  |  |  |  |  |  |
| 0.011<br>(0.00001) | 0.011<br>(0.00001) | 1.0001<br>(0.0001) | 0.9842<br>(0.0004) | 0.007<br>(0.00001) | 0.007<br>(0.00001) | 1.0004<br>(0.00035) | 0.5372<br>(0.0002) |
| 0.011<sup>a</sup> | 0.011<sup>a</sup> | 1.0001<sup>c</sup> |  | 0.010<sup>a</sup> | 0.010<sup>a</sup> | 1.0004<sup>c</sup> |  |
| 0.010<sup>b</sup> | 0.010<sup>b</sup> |  |  | 0.010<sup>b</sup> | 0.010<sup>b</sup> |  |  |
| 0.011<sup>c</sup> | 0.011<sup>c</sup> |  |  | 0.010<sup>c</sup> | 0.010<sup>c</sup> |  |  |
| **C = 0** |  |  |  |  |  |  |  |
| 0.094<br>(0.00004)<br>0.246<sup>a</sup><br>0.505<sup>b</sup><br>0.258<sup>c</sup> | 0.019<br>(0.00001)<br>0.044<sup>a</sup><br>0.091<sup>b</sup><br>0.046<sup>c</sup> | 1.0002<br>(0.0004)<br>1.0003<sup>c</sup> | 0.5533<br>(0.0002)<br>0.5536<br>(0.0002)<sup>d</sup> | 0.105<br>(0.00009)<br>0.190<sup>a</sup><br>0.137<sup>b</sup><br>0.212<sup>c</sup> | 0.077<br>(0.00008)<br>0.097<sup>a</sup><br>0.070<sup>b</sup><br>0.108<sup>c</sup> | 1.0082<br>(0.0007)<br>1.0083<sup>c</sup> | 0.1966<br>(0.0001)<br>0.1956<br>(0.0001)<sup>d</sup> |
| **C = 0.1** |  |  |  |  |  |  |  |
| 0.092<br>(0.00004)<br>0.240<sup>a</sup><br>0.459<sup>b</sup><br>0.251<sup>c</sup> | 0.020<br>(0.00001)<br>0.043<sup>a</sup><br>0.082<sup>b</sup><br>0.043<sup>c</sup> | 1.0005<br>(0.0004)<br>1.0003<sup>c</sup> | 0.5535<br>(0.0002) | 0.096<br>(0.00008)<br>0.161<sup>a</sup><br>0.114<sup>b</sup><br>0.179<sup>a</sup> | 0.070<br>(0.00007)<br>0.082<sup>a</sup><br>0.058<sup>b</sup><br>0.092<sup>c</sup> | 1.0311<br>(0.0007)<br>1.0078<sup>c</sup> | 0.1964<br>(0.0001) |
| **C = 1** |  |  |  |  |  |  |  |
| 0.079<br>(0.00004)<br>0.200<sup>a</sup><br>0.262<sup>b</sup><br>0.210<sup>c</sup> | 0.016<br>(0.00001)<br>0.036<sup>a</sup><br>0.047<sup>b</sup><br>0.037<sup>c</sup> | 1.0008<br>(0.0004)<br>1.0003<sup>c</sup> | 0.5533<br>(0.0002) | 0.071<br>(0.00006)<br>0.091<sup>a</sup><br>0.066<sup>b</sup><br>0.103<sup>c</sup> | 0.035<br>(0.00005)<br>0.047<sup>a</sup><br>0.034<sup>b</sup><br>0.061<sup>c</sup> | 1.0053<br>(0.0007)<br>1.0046<sup>c</sup> | 0.1962<br>(0.0001) |
| **C = 10** |  |  |  |  |  |  |  |
| 0.032<br>(0.00002)<br>0.056<sup>a</sup><br>0.052<sup>b</sup><br>0.056<sup>c</sup> | 0.005<br>(0.00001)<br>0.010<sup>a</sup><br>0.009<sup>b</sup><br>0.011<sup>c</sup> | 1.0004<br>(0.0004)<br>1.0001<sup>c</sup> | 0.5532<br>(0.0002) | 0.008<br>(0.00002)<br>0.016<sup>a</sup><br>0.015<sup>b</sup><br>0.016<sup>c</sup> | 0.005<br>(0.00001)<br>0.008<sup>a</sup><br>0.007<sup>b</sup><br>0.008<sup>c</sup> | 1.0004<br>(0.0007)<br>1.0004<sup>c</sup> | 0.1956<br>(0.0001) |
| **C = 0** |  |  |  |  |  |  |  |
| 0.029<br>(0.00001)<br>0.034<sup>a</sup><br>0.070<sup>b</sup><br>0.036<sup>c</sup> | 0.029<br>(0.00001)<br>0.034<sup>a</sup><br>0.070<sup>b</sup><br>0.036<sup>c</sup> | 1.0002<br>(0.0001)<br>1.0002<sup>c</sup> | 0.9842<br>(0.00004)<br>0.9841<br>(0.00004)<sup>d</sup> | 0.060<br>(0.00004)<br>0.089<sup>a</sup><br>0.064<sup>b</sup><br>0.101<sup>c</sup> | 0.060<br>(0.00004)<br>0.089<sup>a</sup><br>0.064<sup>b</sup><br>0.101<sup>c</sup> | 1.0048<br>(0.0004)<br>1.0054<sup>c</sup> | 0.5387<br>(0.0002)<br>0.5375<br>(0.0002)<sup>d</sup> |
| **C = 0.1** |  |  |  |  |  |  |  |
| 0.028<br>(0.00001)<br>0.033<sup>a</sup><br>0.064<sup>b</sup><br>0.035<sup>c</sup> | 0.028<br>(0.00001)<br>0.033<sup>a</sup><br>0.064<sup>b</sup><br>0.035<sup>c</sup> | 1.0002<br>(0.0001)<br>1.0002<sup>c</sup> | 0.9842<br>(0.00004) | 0.055<br>(0.00003)<br>0.082<sup>a</sup><br>0.058<sup>b</sup><br>0.093<sup>a</sup> | 0.055<br>(0.00003)<br>0.082<sup>a</sup><br>0.058<sup>b</sup> | 1.0043<br>(0.0004)<br>1.0051<sup>c</sup> | 0.5386<br>(0.0002) |
| **C = 1** |  |  |  |  |  |  |  |
| 0.023<br>(0.00001)<br>0.027<sup>a</sup><br>0.036<sup>b</sup><br>0.028<sup>c</sup> | 0.023<br>(0.00001)<br>0.027<sup>a</sup><br>0.036<sup>b</sup><br>0.028<sup>c</sup> | 1.0001<br>(0.0001)<br>1.0002<sup>c</sup> | 0.9842<br>(0.0004) | 0.028<br>(0.00002)<br>0.045<sup>a</sup><br>0.032<sup>b</sup><br>0.051<sup>a</sup> | 0.028<br>(0.00002)<br>0.045<sup>a</sup><br>0.032<sup>b</sup><br>0.051<sup>a</sup> | 1.0024<br>(0.0004)<br>1.0030<sup>c</sup> | 0.5380<br>(0.0002) |
| **C = 10** |  |  |  |  |  |  |  |
| 0.007<br>(0.00001)<br>0.007<sup>a</sup><br>0.007<sup>b</sup><br>0.007<sup>c</sup> | 0.007<br>(0.00001)<br>0.007<sup>a</sup><br>0.007<sup>b</sup><br>0.007<sup>c</sup> | 1.0001<br>(0.0001)<br>1.0001<sup>c</sup> | 0.9842<br>(0.0004) | 0.004<br>(0.00001)<br>0.007<sup>a</sup><br>0.006<sup>b</sup><br>0.007<sup>c</sup> | 0.004<br>(0.00001)<br>0.007<sup>a</sup><br>0.006<sup>b</sup><br>0.007<sup>c</sup> | 1.0007<br>(0.0003)<br>1.0002<sup>c</sup> | 0.5373<br>(0.0002) |
| **C = 0** |  |  |  |  |  |  |  |
| 0.062<br>(0.00003)<br>0.165<sup>a</sup><br>0.034<sup>b</sup><br>0.171<sup>c</sup> | 0.013<br>(0.00001)<br>0.030<sup>a</sup><br>0.061<sup>b</sup><br>0.031<sup>c</sup> | 1.0023<br>(0.0004)<br>1.0002<sup>c</sup> | 0.5534<br>(0.0002)<br>0.5536<br>(0.0002)<sup>d</sup> | 0.069<br>(0.00006)<br>0.121<sup>a</sup><br>0.087<sup>b</sup><br>0.137<sup>c</sup> | 0.051<br>(0.00006)<br>0.062<sup>a</sup><br>0.044<sup>b</sup><br>0.070<sup>c</sup> | 1.0056<br>(0.0007)<br>1.0054<sup>c</sup> | 0.1962<br>(0.0001)<br>0.1956<br>(0.0001)<sup>d</sup> |
| **C = 0.1** |  |  |  |  |  |  |  |
| 0.061<br>(0.00003)<br>0.161<sup>a</sup><br>0.309<sup>b</sup><br>0.178<sup>c</sup> | 0.013<br>(0.00001)<br>0.029<sup>a</sup><br>0.055<sup>b</sup><br>0.032<sup>c</sup> | 1.0024<br>(0.0004)<br>1.0002<sup>c</sup> | 0.5531<br>(0.0002) | 0.063<br>(0.00006)<br>0.111<sup>a</sup><br>0.079<sup>b</sup><br>0.178<sup>c</sup> | 0.046<br>(0.00005)<br>0.057<sup>a</sup><br>0.040<sup>b</sup><br>0.091<sup>c</sup> | 1.0198<br>(0.0007)<br>1.0051<sup>c</sup> | 0.1961<br>(0.0001) |
| **C = 1** |  |  |  |  |  |  |  |
| 0.053<br>(0.00003)<br>0.131<sup>a</sup><br>0.172<sup>b</sup><br>0.137<sup>c</sup> | 0.010<br>(0.00001)<br>0.024<sup>a</sup><br>0.031<sup>b</sup><br>0.025<sup>c</sup> | 1.0010<br>(0.0004)<br>1.0002<sup>c</sup> | 0.5532<br>(0.0002) | 0.033<br>(0.00004)<br>0.061<sup>a</sup><br>0.044<sup>b</sup><br>0.070<sup>c</sup> | 0.023<br>(0.00003)<br>0.031<sup>a</sup><br>0.023<sup>b</sup><br>0.035<sup>c</sup> | 1.0021<br>(0.0007)<br>1.0030<sup>c</sup> | 0.1959<br>(0.0001) |
| **C = 10** |  |  |  |  |  |  |  |
| 0.021<br>(0.00001)<br>0.035<sup>a</sup><br>0.033<sup>b</sup><br>0.035<sup>c</sup> | 0.003<br>(0.00001)<br>0.006<sup>a</sup><br>0.006<sup>b</sup><br>0.006<sup>c</sup> | 1.0000<br>(0.0004)<br>1.0001<sup>c</sup> | 0.5531<br>(0.0002) | 0.006<br>(0.00001)<br>0.009<sup>a</sup><br>0.008<sup>b</sup><br>0.010<sup>c</sup> | 0.003<br>(0.00001)<br>0.005<sup>a</sup><br>0.004<sup>b</sup><br>0.006<sup>c</sup> | 0.9997<br>(0.0007)<br>1.0002<sup>c</sup> | 0.1953<br>(0.0001) |

An approximate treatment can be applied for the case when *S = 2Ns* and *T = 2Nt* are both < 1, on the same lines as for Equations 15-20 for the mutation-selection balance case. The coefficients for the approximate recursion relations for *Y*_4_ and *Y*_5_ are as follows:

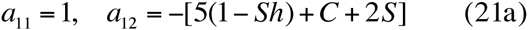

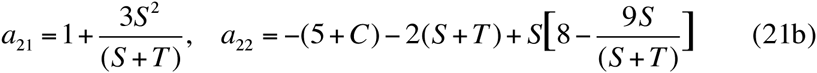

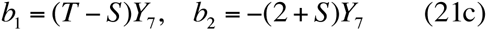

The equivalents of Equations 17 and 18 are given in section 3 of the Supplementary Information, which can be used to calculate ∆*H*_*sel*_ in the same way as for the mutation-selection case. The first-order term in *S* and *T* in ∆*H*_*sel*_ gives:

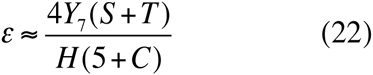

The inbred load in this case is (*s + t*)E{*pq*} (see Equations 9), so that this is identical in form to the first-order approximation for the mutation-selection balance case (see Equation 19). For sufficiently weak selection, therefore, heterozygote advantage should always lead to an enhancement of neutral variability at a linked locus.

We note, however, that the asymptotic rate of loss of variability at a locus with heterozygote advantage has a complex dependence on the deterministic equilibrium allele frequency and the population size, with extreme allele frequencies (outside the range 0.2 to 0.8) leading to an acceleration of loss of variability when *S + T* is sufficiently large (Robertson 1962). This suggests that a similarly complex pattern should be found for the dependence of the measure of retardation/acceleration based on the leading eigenvalue of the recursion matrix. Figure S3 of the Supplementary Information (section 9) shows that this is indeed the case, although there is only a small region of parameter space where acceleration rather than retardation of loss of variability occurs.

Under most circumstances, some retardation of the loss of variability at a neutral locus is, therefore, likely to be observed as a result of linkage to a selected locus, although its magnitude is small unless linkage is tight and *S + T* ≫ 1. For *C* ≫ *S + T* it is unlikely that any experimentally detectable effects could be observed. The same applies to the apparent selection coefficients against homozygotes at the neutral locus.

## Discussion

### Some general considerations

Our results shed some new light on the old question of the existence and properties of associative overdominance (AOD) at a neutral locus in a randomly mating population of finite size, resulting from randomly generated linkage disequilibrium with respect to a locus subject to selection. The pioneering work of Sved (1968, 1971, 1972), Ohta and Kimura (1970) and Ohta (1971) gave expressions for the apparent selection coefficients against homozygotes at a neutral locus, using the equivalents of our Equations 3 and 8. Our approximate estimates of the apparent selection coefficients for the case of a finite population founded from an infinite population at equilibrium with no LD between the selected and neutral loci (Equations 4 and 9) followed their approach in using only the terms involving *D*^2^ in these equations, which are the dominant terms.

However, as was noted by Sved (1968, p.552) for the case of heterozygote advantage and by Latter (1998) for the case of selection against deleterious mutations, these selection coefficients do not result in any change in allele frequency at the neutral locus. As mentioned above (Equation 10), this is a completely general conclusion: application of the Price equation (Price 1970) shows that the change **A***x* per generation in the frequency of allele A_1_ at the neutral locus is equal to *aD*, where *a* is the average effect of A_1_ on fitness (see also Santiago and Caballero 1995, p.1016), to the order of the approximations used here (neglect of second-order terms in *s* and 1/*N)*. Neither the change in allele frequency nor the change in heterozygosity at the neutral locus are influenced by terms in *D*^2^.

There is thus no connection between apparent overdominance caused by linkage disequilibrium and any retardation of loss of variability by drift at a neutral locus. Such a connection seems to have been widely assumed to be the basis for a retardation of loss of variability (e.g. Latter 1998), although it had been pointed out by Charlesworth (1991) that AOD caused by identity disequilibrium has no effect on allele frequencies at the neutral loci concerned. As was noted by Bierne *et al*. (2000), for this case it is necessary to treat apparent overdominance at the neutral locus as a phenomenon that is entirely distinct from any retardation of loss of variability. This point is reinforced by the results in Tables 1 and 2 for the case of selection against recurrent deleterious mutations, which show that apparent overdominance can accompany an acceleration of loss of neutral variability when the dominance coefficient *h* is sufficiently large (see the entries with *h* = 0.45). Of course, *h* < ½ is a necessary condition for both retardation of loss of variability and for apparent overdominance. This is not restrictive, given the evidence that the dominance coefficients for slightly deleterious mutations are generally < ½ (Crow 1993; Manna *et al*. 2011).

### Selection against deleterious mutations

Computer simulations of systems with many loci subject to mutation to deleterious alleles have shown that a noticeable retardation of loss of neutral variation at linked loci can occur (Latter 1998; Pamilo and Palsson 1998, 1999; Wang and Hill 1999; Wang *et al*. 1999), although only for certain ranges of values of selection and dominance coefficients. As pointed out by Pamilo and Palsson (1998, 1999), Wang and Hill (1999) and Wang *et al*. (1999), there is a conflict between the effect of AOD on neutral variability and the effect of background selection (BGS); the latter involves a reduction in the effective population size experienced by a neutral locus caused by its linkage to deleterious mutations (Charlesworth *et al*. 1993). Their multi-locus simulations, as well as the matrix-based investigation of a model of a single selected locus linked to a neutral locus sib-mating and selfing lines by Wang and Hill (1999), showed that retardation versus acceleration of the rate of loss of neutral variability is favoured by relatively weak selection (*2Ns* values of order four or less) and low dominance coefficients; see, for example, Figure 1 of Wang and Hill (1999).

Our analytical and numerical methods, employing a 9 x 9 matrix of ‘moments’ of functions of the allele frequencies at the two loci and *D*, as well as the weak selection approximations for the change in heterozygosity, have allowed us to investigate in detail the regimes in which retardation versus acceleration of loss of neutral variability occurs.

The main conclusions are that:

(1) Retardation rather than acceleration of loss of variability is favored by sufficiently low values of the dominance coefficient, *h*, when *S = 2Ns* is in the range 0.5 to 4.5 (Figure 3). The commonly used estimate of *h* = 0.25 for slightly deleterious mutations (Manna *et al*. 2011; Charlesworth 2015) is consistent with retardation when *S* ≤ 2.5, except when *q\** is close to 0.5.
(2) *S < 0.5* allows retardation of loss even for *h* values approaching ½; this domain of *S* values can be thought of as the *AOD limit*.
(3) In contrast, there is always an acceleration of loss of variability when *S* > 4, even with very low *h* values; this constitutes the *BGS limit*. Despite this acceleration, apparent heterozygote advantage at the neutral locus is always observed, provided that *h* < ½.
(4) For *q\** ≪ 1and for *S* ≫ 1, the asymptotic behaviour of the expected heterozygosity at the neutral locus is well predicted by the *N*_*e*_ value given by the standard equation for BGS.
(4) The magnitude of the extent of retardation or acceleration of loss of variability is an increasing function *of C/S*, for a given *h* value. Equation (19) also shows that, when *S ≪* 1, the effect is proportional to the inbred load scaled by *2N*, for a given value of *C*.

Population genomic analyses of levels of nonsynonymous site variability in a variety of organisms have consistently shown that there is a wide distribution of selection coefficients against new deleterious mutations, and suggest a mean *S* for natural populations that is ≫ 1 (reviewed by Charlesworth 2015). The large effective population sizes of most populations that have been studied in this way imply, however, that the mean selection coefficient against a new deleterious mutation is likely to be very small. For example, the current estimate of the mean *hs* for new nonsynonymous mutations in *D. melanogaster* is approximately 0.001; with *h* = 0.25, the corresponding mean *s* is 0.004 (Charlesworth 2015). With *N* = 50, as used in the examples below, the mean *S* is 0.4. If there were no variation in *s* for new mutations, this would suggest that they fall well within the range where Equations 18 and 19 predict retardation of loss of variability. However, as discussed below we also have to take into account the fact that there is a wide distribution of *s*. A coefficient of variation for *s* for new mutations of approximately 2 is suggested by the Drosophila polymorphism data (Kousathanas and Keightley 2013).

There is also evidence for deleterious mutations with larger fitness effects than typical nonsynonymous mutations, such as insertions and deletions; these appear to make up only a small contribution to the spectrum of new mutations (Charlesworth 2015), and will be neglected in what follows, since they must be sparsely distributed across the genome. Their net effect is probably to tilt the balance slightly more in favor of BGS than if they are ignored (but see the section below on relevant data).

### Effects of multiple loci subject to mutation and selection

We now consider how the effects of multiple loci subject to deleterious mutation affect a linked neutral locus; this question can asked about both the apparent selection coefficients and the extent of retardation or acceleration of loss of variability.

#### Apparent selection coefficients

With respect to the apparent selection coefficients, previous workers (Ohta 1971, 1973; Ohta and Cockerham 1974; Bierne *et al*. 2000) assumed that multiple loci combine approximately additively. Table 2, which describes results for a pair of selected loci with, suggests that this assumption may be somewhat conservative; in most of the examples shown there, the selection coefficients are somewhat larger than twice the corresponding single locus values, especially with a low initial frequency of the A_1_ allele.

The procedure of summing contributions over all sites when there are many selected loci, as used by Ohta (1971, 1973), may thus underestimate the apparent selection coefficients, although the departure from additivity is likely to be small for the very weak selection that is probably most common. An alternative to Equation 13 of Ohta (1973), based on Equations 3, is derived in section 6 of the Supplementary Information (Equations S31). We assume that the population is being studied at a sufficiently long time after establishment that its LD is close to equilibrium. We also assume a single chromosome with a uniform rate of recombination along the chromosome, as well as a linear genetic mapping function (this somewhat overestimates the amount of recombination, compared with the true situation with partial crossover interference). For a randomly placed marker, we have the asymptotic result:

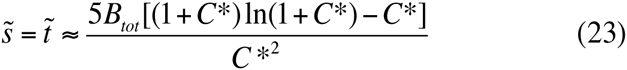

where *B*_*tot*_ is the inbred load associated with homozygosity for the whole schromosome, and *C\** = *2NM*, where *M* is the effective map length of the chromosome (taking into account any sex differences in recombination rates).

It is useful to compare this formula with the multi-locus simulation results of Latter (1998), who modeled a *D. melanogaster* autosome evolving in a population maintained at a size of 50 for 200 generations after its foundation from an equilibrium population. The chromosome was 100cM in length; the absence of crossing over in males means that the effective value of *M* is 0.5 Morgans, so that *C*\* = 50 with *N* = 50. For a randomly placed marker, Equation 23 gives an approximate apparent selection coefficient of *0.312B*_*tot*_.

*B*_*tot*_ can be estimated as follows. The initial value of the net fitness (relative to a balancer chromosome) for *D. melanogaster* second chromosomes that were purged of major effect mutations by extracting them from a population maintained for many generations at an approximate size of 50 was 0.4 (Latter *et al*. 1995; Latter 1998). This is equivalent to the fitness of a homozygous chromosome relative to that of an outbred genotype in the initial population; the value of *B*_*tot*_ for the initial population is then - ln(4) = 0.90, assuming multiplicative fitness effects. This assumption is consistent with the log-linear decline in fitness with inbreeding coefficient *F* in the small laboratory populations of *D. melanogaster* described by Latter *et al*. (1995).

The simulations of Latter (1998) adjusted the number of mutable loci to produce an initial *B*_*tot*_ of 0.90. However, we need to take into account the reduction in *B*_*tot*_ as variability is lost at the selected loci. A minimum estimate of *B*_*tot*_ at generation *t* is provided by assuming that the ratio of E{*pq*} to its initial value follows the standard neutral result, 1 - *F*_*t*_ = *exp(-t/2N)*. For *N* = 50 and *t* =200, this procedure yields *B*_*tot*_ = 0.90 x (1 - *F*_200_) = 0.122, and a value of 0.038 for the apparent selection coefficient on a random marker, somewhat lower than the value of 0.05 given in Figure 6 of Latter (1998). The discrepancy probably reflects an overestimate of the loss of variability by use of the standard neutral value of *F*_*t*_. Latter’s Figure 4 shows *F*_200_ = 0.7 for the selected loci in his simulations, giving *B*_*tot*_ = 0.270 and an apparent selection coefficient of 0.084. The single locus approximations for the apparent selection coefficients often overestimate the true values for later generations (see Table 1), so that it is not surprising that Equation 23 overestimates the apparent selection coefficient.

Since the numerator of Equation (23) is only a logarithmic function of *C\**, the apparent selection coefficients are roughly inversely proportional to population size. Species with larger numbers of chromosomes, and with crossing over in both sexes, will have much smaller apparent selection coefficients for the same value of *N*. Detectable apparent selection coefficients arising from AOD in randomly mating populations are thus only likely to be found in very small populations, especially in organisms with large numbers of chromosomes such as most vertebrates. This suggests that AOD arising from identity disequilibrium (Szulkin *et al*. 2010) is a more likely explanation of heterozygosity/fitness relations in natural populations than LD due to drift in a randomly mating population (Hansson and Westerberg 2002), unless the population has been reduced to a very small number of breeding individuals.

#### Rate of loss of variability

This approach can also be applied to the rate of loss of neutral variability, using the weak selection approximation of Equation 19, which includes only the first and second-order terms in *S* in the expression for ε = *N*_*e*_ /*N* - 1 = 2*N* ∆*H*_*sel*_, again assuming that LD is close to neutral equilibrium. The derivation of the approximate expected value of *ε* for a neutral marker placed randomly on a chromosome, using the same assumptions as before, is given in section S7 of the Supplementary Information (Equations S32-S35). The final result is:

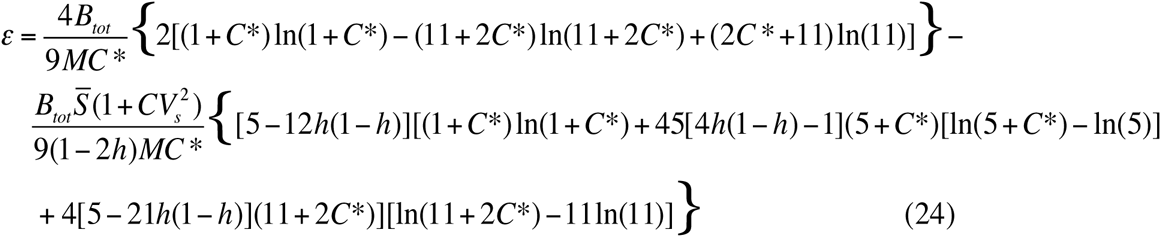

where *CV*_*s*_ is the coefficient of variation of the distribution of *s* for new mutations. With an exponential distribution, *CV*_*s*_ = 1.

We can compare the predictions of this equation with the simulation results in Latter’s Table 6 for a mean *S* of 0.5, which is similar to the value suggested by the population genomic analyses for a population size of 50 and meets the weak selection requirement. Latter used a measure (∆*F*) derived from net change in *F* for neutral loci over *t* generations, relative to the corresponding value without selection; this is approximately equivalent to the ratio of *N* to the harmonic mean of *N*_*e*_, i.e., to 1/(1 + *ε*). For the case of an exponential distribution of *s*, he estimated ∆*F* as approximately 0.6 for *t* = 200, for three different dominance coefficients, 0, 0.1 and 0.2. With *h* = 0.2, the estimate of *ε* for a random marker with a population size of 50 is 1.35*B*_*tot*_; with the above estimate of *B*_*tot*_ = 0.270 for generation 200, this gives *ε* = 0.365 and ∆*F* = 0.73, which is somewhat higher than Latter’s value. But the comparisons of the single locus simulation results with Equation 19 suggested that the approximation underestimates *ε* by about 30% (Figure 4); increasing *ε* by this amount to 0.474 give ∆*F* = 0.68. If the effects of different loci on *N*_*e*_ combine multiplicatively rather than additively, as is known to be the case for BGS (Charlesworth and Charlesworth 2010, p.402), *ε* should be replaced by exp(ε) - 1 = 0.606, and ∆*F* = 0.62, which is very close to the simulation value. The value of *h* has only a small effect on the results; for example, with *h* = 0, the uncorrected *ε* = 0.454 and ∆*F* = 0.69.

We can also use Equation 24 to determine the expectations with the population genomics estimates of mean *S* of 0.004 and *CV*_*s*_ = 2, with *h* = 0.25. Substituting these values into Equation 24 with the same *N* and recombination parameters as before, Equation 24 gives a negative values of *ε* = −0.0663, and ∆*F* = 1.07. This implies that BGS would cause a reduction in *N*_*e*_, to about 93% of the neutral value. This conclusion should be treated with caution, since the very wide distribution of selection coefficients in this case means that there will be a substantial range of values that violate the assumption of *S* < 1 required for the validity of Equation 19.

### Relation to the data on loss of variability in small populations

Most of the data that are relevant to the question of whether AOD generated by linkage to deleterious mutations can retard the loss of neutral variability in small randomly mating populations comes from a few studies of the behavior of putatively neutral markers and quantitative traits in small laboratory populations of *D. melanogaster*. In order to avoid the complications of interpretation associated with high levels of inbreeding, we will not discuss results from studies of sib or first-cousin mating, although these suggest a significant retardation of loss of variability (Rumball *et al*. 1994), which is consistent with the conclusion that retardation is favored when *S* is small. Latter (1998) analyzed the results of Latter *et al*. (1995) for two allozyme markers, and found a ∆*F* value of 0.52 over 200 generations in populations maintained at a size of approximately 50 (see his Table 6), consistent with his multi-locus simulation results based on an exponential distribution of *s*.

Experiments using seven allozyme markers and two bristle traits in pedigreed populations ranging in size from sib-mating to 500, and maintained for 50 generations, showed that the regression coefficient of heterozygosity and genetic variance on pedigree inbreeding coefficient were significantly lower than the value of 1 expected with neutrality, suggesting a retardation of loss of variability (Gilligan *et al*. 2005). The facts that the results from populations with different sizes were pooled, and that there is non-linear relation between heterozygosity and inbreeding coefficient when AOD is acting, mean that a quantitative analysis of these results in terms of our model is impossible. In addition, a later analysis of 8 microsatellite loci over 48 generations for the same set of populations showed an acceleration of loss of variability by this criterion (Montgomery *et al*. 2010), which the authors interpreted as evidence for selective sweeps related to adaptation to the laboratory environment.

Different experiments have, therefore, given very different patterns, and it is certainly possible that any retardation of loss of variability due to AOD may be obscured by selective sweeps even if *N is* of the size that would otherwise lead to a retardation of loss of variability. However, a basic problem with experiments of this type is that the heterozygosity for a single neutral locus has a very high stochastic variance, even if allele frequencies are estimated with complete accuracy (Avery and Hill 1977); see Equation S39 of the Supplementary Information, section 8. This yields the asymptotic expression for large *t*:

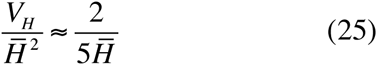

This equation implies that the relative error increases as mean heterozygosity decreases. For example, at generation 100, if the initial heterozygosity was at the maximal value of 0.5 for a biallelic locus, the mean neutral *H* for *N* = 50 is 0.184, yielding an expected coefficient of variation for a single locus in a single population of 2.17. To obtain a coefficient of variation of 0.1, (2.17/0.1)^2^ = 471 independent heterozygosities would need to be measured, either from independent loci or populations, or from a combination of the two. Given that Figure 6 of Latter (1998) shows a difference of approximately 0.2 between the expected neutral *F* value and the value from simulations with AOD due to deleterious mutations, which corresponds to the difference in scaled *H* values, it is clear that many more replications than this would be necessary to obtain statistically significant results, especially as the distribution of *H* is far from normal. The differences between the different experiments could thus be purely stochastic, especially as the autocorrelations between *H* values in different generations means that the regression tests used by Gilligan *et al*. (2005) and Montgomery *et al*. (2010) are problematic.

It is possible that tight linkage of neutral markers to sets of very strongly and highly recessive deleterious variants that are in repulsion LD with each other could maintain variation as a result of the pseudo-overdominance that can arise in this circumstance: see the section on two selected loci above, and Charlesworth and Charlesworth (1997), Pamilo and Palsson (1999) and Palsson (2001). This is most likely to occur in very small populations; major effect mutations are relatively sparsely distributed, so that their effects on neutral variability are likely to be restricted to specific genomic regions, rather than evenly spread across the genome. Recent genomic investigations of the well-known Chillingham population of cattle, which have been maintained at very small population size for 350 years, are consistent with this interpretation. Genome-wide diversity is very low, and residual variability is localized to a number of specific genomic locations (Willams *et al*. 2016).

A full assessment of the possibility that a realistic distribution of mutational effects on fitness (DFE), which are currently becoming available from genome-wide polymorphism data (Charlesworth 2015), can cause significant retardation of the loss of neutral variability will require simulations that use realistic DFEs, coupled with experiments on the effects of reduced population size that exploit the high levels of replication that can be achieved using modern genomic technology for generating large numbers of markers. It would also be desirable to use populations that have been maintained for a long period in the laboratory at a large size as the initial population, to avoid possible confounding effects of selective sweeps. Current theory and data cannot convincingly answer the question of whether AOD due to deleterious mutations is a credible explanation for the presence of more than expected levels of variability in small populations.

### Selection in favor of heterozygotes

Genome scans for signatures of balancing selection suggest that this is a relatively rare phenomenon compared with the number of genes in the genome (Charlesworth 2006; Gao *et al*. 2015), so that it is unlikely that a given neutral site will be closely linked to a locus maintained by heterozygote advantage. The exception is inversion polymorphisms in organisms such as Drosophila; these can cover relatively large proportions of the genome, and also suppress crossing over to a considerable extent outside their breakpoints (Krimbas and Powell 1992). They could thus have a substantial impact on the behavior of neutral variants in small populations. Thus, our conclusions concerning heterozygote advantage are relevant to experimental results on small populations where inversions are segregating, as is often the case with *D. melanogaster*.

Our main conclusion is that retardation of loss of variability is nearly always observed when there is heterozygote advantage, except when the allele frequency at the selected locus is close to the boundaries 0 or 1, and there is relatively strong selection, when it is known that heterozygote advantage tends to accelerate the loss of variability (Robertson 1962). As shown in Figure 4 and Table 3, a substantial degree of retardation can occur when *c* ≪ 10(*s + t*), especially with intermediate equilibrium allele frequencies. As with mutation and selection, with *S + T* < 1 and a fixed value of *C* the magnitude of retardation is proportional to the inbred load scaled by 2*N* (see Equation 22).

## Acknowledgments

We thank Nick Barton and Bill Hill for useful discussions. Lei Zhao was supported by a China Scholarship Council (CSC) scholarship (#201506100068).

## Appendix

### Approximations for the expected frequency of B_2_ and expected heterozygosity at the selected locus

With weak selection, the deterministic change in the frequency *q* of allele B_2_ under selection and mutation (neglecting terms of order *s*^2^) is given by:

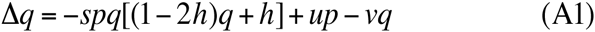

Because ∆*q* is third degree in *q*, it is impossible to obtain exact, closed equations for the changes in the expectations of *q* and *q*_2_, E{*q*}, and E{*q*^2^} under drift, selection and mutation. However, if it is assumed that second-order terms in the deviation of *q* from its deterministic equilibrium value *q\**, given by Equation 5a, can be ignored, approximate recursion relations can be obtained as follows. (The matrix calculations described in section 1 of the Supplementary Information, which neglect only third-order terms in (*q - q\**), yield better approximations.)

Taking the first derivative of Equation A1 with respect to *q* yields a linear deterministic recursion relation for the departure of *q* in generation *t* from the equilibrium frequency *q\** under mutation and selection, which is given by Equations 5:

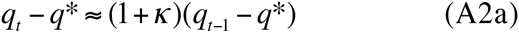

where

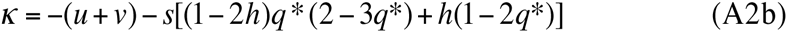

Since this is linear in *q*, the mean of *q* under drift, selection and mutation remains at *q\**.

The equilibrium expected heterozygosity at the selected locus is given by *H*_2_ = 2*q*\*(1 - *q*\*)*A*/(1+*A*), where *A* = −4*N*_*e*_κ (e.g., Charlesworth and Charlesworth 2010, p.355). The change in the expected heterozygosity at the selected locus at time *t, H*_2*t*_, is approximately equal to the expectation of 2(1 - 2*q*_*t*_)∆*q*(*q*_*t*_); combining this with Equations A2 yields the following recursion relation:

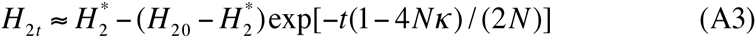

Because *H*_2_ is twice the expectation of *pq*, Equation A3 was used in Equations 4 for determining the approximate apparent selection coefficients.

### Approximate neutral recursion relations for the expectation of *r*^2^

For the purely neutral case, Sved (1972) proposed the following approximation for the recursion relation for the expected correlation coefficient in generation *t*, E{*r*_*t*_^2^}:

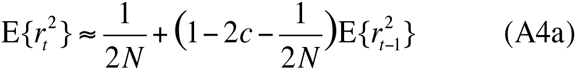

where *c* is the frequency of recombination between the two loci, assumed to be ≪ 0.5. Since mutation at the B locus reduces the coefficient of linkage disequilibrium *D* with the A locus by *u + v* each generation, this equation should be modified as follows:

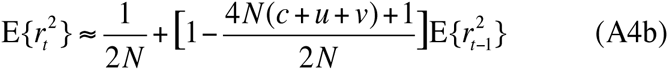

This yields the equilibrium solution:

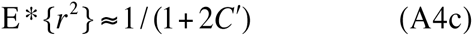

where *C’* = 2*N(c + u + v)*.

If the initial population is in linkage equilibrium, Equations A4 yield the following homogeneous recursion relation:

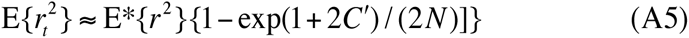

This suggests the following expression for σ_*d*_^2^ = E{*D*^2^}/E{*xypq*}:

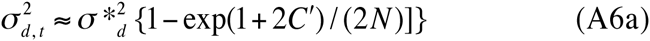

where the equilibrium value of σ_*d*_^2^ is given by the following expression (Ohta and Kimura 1971):

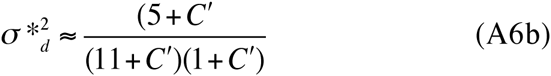

The simulation results show that Equation A6a tends to overestimate E{*r*_2_} up to *t* = 0.5*N*; it is quite accurate for later generations up to *t = N*, and tends to underestimate E{*r*_2_} thereafter. Equation A5 gives a better approximation for the earlier and later generations, but a worse approximation for the intermediate generations.

### Expectations of the reciprocals of the allele frequencies at the neutral locus

The expectations of *x*^−1^ and *y*^−1^ = (1 - *x*)^−1^, conditioned on segregation at locus A, for use in Equations 3 and 9, can be derived using the diffusion equation solution for the case of pure drift (Kimura 1955), under the assumption that departures from strict neutrality can be neglected as a first-order approximation. For this purpose, it is convenient to rescale time in units of 2*N* generations, such that *T = t/(2N)*. Using the first three terms in the series expansion for the unconditional probability density of the frequency *x* of A_1_at time *T*, given an initial frequency of *x*_0_, we have:

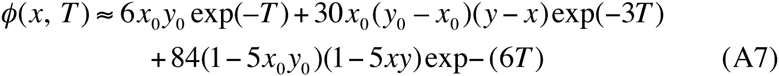

The expected frequency of heterozygotes at locus A at time *T* is

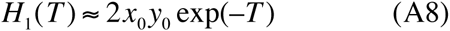

To determine the conditional probability of *x* for a segregating population, we need the probability that the population is still segregating for A_1_ and A_2_ at time *T*, given by Equation 8.4.9 of Crow and Kimura (1970). Using the first two terms in the series, we have

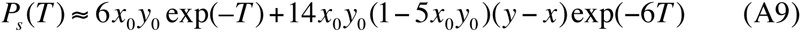

The conditional probability density of *x* at time *T* is then given by φ/*P*_*s*_, which can be used to determine the mean of *x*^−1^ at time *T* for segregating populations:

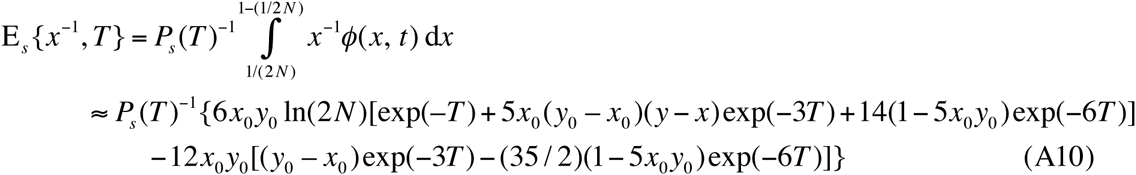

An equivalent expression for the expectation of *y*^−1^ can be obtained by interchanging *x* and *y* and *x*_0_ and *y*_0_ in these equations.

For sufficiently large *T*, Equation A7 implies that the conditional probability distribution becomes uniform, regardless of the initial frequency, so that its mean should be well approximated by

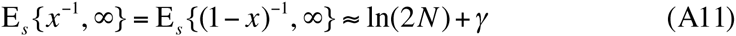

where *γ* is Euler’s constant (approximately 0.5772) for the difference between the sum and integral of 1/*x*.

As was pointed out by Fisher (1930), there is a slight departure of the asymptotic distribution from a uniform distribution, due to the inaccuracy of the diffusion approximation near the frequencies 0 and 1, with fewer frequencies in the subterminal classes close to either boundary, although the distribution is still symmetrical about 0.5. Using the table of exact values computed by Fisher for the first 10 classes from 1/(*2N*) upwards, and from 1 - 1/(*2N*) downwards, the asymptotic conditional means are slightly greater than the above value, by 2.299 - [2.6556 x *2N/(2N* - 1)]. Addition of this term to Equation A11 provides a slightly more accurate approximation for the long-term expectation of the reciprocals of the allele frequencies, which was used to calculate the asymptotic expected selection coefficients.

## Literature Cited

Avery, P. J., and W. G. Hill, 1977 Variability in genetic parameters among small populations. Genet. Res. 29: 193–213.

Balick, D. J., R. Do, C. A. Cassa, D. Reich and S. Sunyaev, 2015 Dominance of deleterious alleles controls the response to a population bottleneck. PloS Genet. 11: e1005436.

Bierne, N., A. Tsitrone and P. David, 2000 An inbreeding model of associative overdominance during a population bottleneck. Genetics 155: 1981–1990.

Charlesworth, B., 2015 Causes of natural variation in fitness: evidence from studies of Drosophila populations. Proc. Natl. Acad. Sci. USA 12: 1662–1669.

Charlesworth, B., and J. L. Campos, 2014 The relations between recombination rate and patterns of molecular evolution and variation in *Drosophila*. Ann. Rev. Genet. 48: 383–403.

Charlesworth, B., and D. Charlesworth, 1997 Rapid fixation of deleterious alleles by Muller’s ratchet. Genet. Res. 70: 63–73.

Charlesworth, B., and D. Charlesworth, 2010 Elements of Evolutionary Genetics. Roberts and Company, Greenwood Village, CO.

Charlesworth, B., M. T. Morgan and D. Charlesworth, 1993 The effect of deleterious mutations on neutral molecular variation. Genetics 134: 1289–1303.

Charlesworth, D., 1991 The apparent selection on neutral marker loci in partially inbreeding populations. Genet. Res. 57: 159–175.

Charlesworth, D., 2006 Balancing selection and its effects on sequences in nearby genome regions. PLoS Genet. 2: e2.

Cockerham, C. C., and B. S. Weir, 1968 Sib-mating with two loci. Genetics 60: 629–640.

Crow, J. F., 1993 Mutation, mean fitness, and genetic load. Oxf Surv. Evol. Biol. 9:3–42.

Crow, J. F., and M. Kimura, 1970 An Introduction to Population Genetics Theory. Harper and Row, New York.

Cutter, A. D., and B. A. Payseur, 2013 Genomic signatures of selection at linked sites: unifying the disparity among species. Nature Rev. Genet. 14: 262–272.

David, P., 1998 Heterozygosity-fitness correlations: new perspectives on old problems. Heredity 80: 531–537.

Ewens, W. J., 2004 Mathematical Population Genetics. 1. Theoretical Introduction. Springer, New York.

Fisher, R. A., 1930 The distribution of gene ratios for rare mutations. Proc. Roy. Soc. Edinburgh 50: 205–220.

Frydenberg, O., 1963 Population studies of a lethal mutant in Drosophila melanogaster. I. Behaviour in populations with discrete generations. Hereditas 50: 89–116.

Gao, Z., M. Przeworski and G. Sella, 2015 Footprints of ancient-balanced polymorphisms in genetic variation: data from closely related species. Evolution 69: 431–466.

Gilligan, D. M., D. A. Briscoe and R. Frankham, 2005 Comparative losses of quantitative and molecular genetic variation in finite populations of Drosophila melanogaster. Genet. Res. 85: 47–77.

Glémin, S., J. Ronfort and T. Bataillon, 2003 Patterns of inbreeding depression and architecture of the load in subdivided populations. Genetics 165: 2193–2212.

Greenberg, R., and J. F. Crow, 1960 A comparison of the effect of lethal and detrimental chromosomes from Drosophila populations. Genetics 45: 1153–1168.

Haddrill, P. R., L. Loewe and B. Charlesworth, 2010 Estimating the parameters of selection on nonsynonymous mutations in Drosophila pseudoobscura and D. miranda. Genetics 185: 1381–1396.

Haldane, J. B. S., 1927 A mathematical theory of natural and artificial selection. Part V. Selection and mutation. Proc. Camb. Phil. Soc. 23: 838–844.

Haldane, J. B. S., 1949 The association of characters as a result of inbreeding. Ann. Eugen. 15: 15–23.

Hansson, B., and L. Westerberg, 2002 On the correlation between heterozygosity and fitness in natural populations. Mol. Ecol. 11: 2467–2474.

Hill, W. G., 2010 Understanding and using quantitative genetic variation. Phil. Trans. R. Soc. B 365: 73–85.

Hill, W. G., and A. Robertson, 1968 Linkage disequilibrium in finite populations. Theor. Appl. Genet. 38.

Hoffman, J. I., F. Simpson, P. David, Rijks, J., Kuiken, T., et al., 2014 High-throughput sequencing reveals inbreeding depression in a natural population. Proc. Natl. Acad. Sci. USA 10: 3775–3780.

Johnson, T., and N. H. Barton, 2005 Theoretical models of selection and mutation on quantitative traits. Phil. Trans. R. Soc. B 360: 1411–1425.

Kimura, M., 1955 Solution of a process of random genetic drift with a continuous model. Proc. Natl. Acad. Sci. USA 41: 144–150.

Kousathanas, A., and P. D. Keightley, 2013 A comparison of models to infer the distribution of fitness effects of new mutations. Genetics 193: 1197–1208.

Krimbas, C. B., and J. R. Powell (Editors), 1992 Drosophila Inversion Polymorphism. CRC Press, Boca Raton, FL.

Latter, B. D. H., 1998 Mutant alleles of small effect are primarily responsible for the loss of fitness with slow inbreeding in *Drosophila melanogaster*. Genetics 148: 1143–1158.

Latter, B. D. H., J. C. Mulley, D. Reid and L. Pascoe, 1995 Reduced genetic load revealed by slow inbreeding in *Drosophila melanogaster*. Genetics 139: 287–297.

Manna, F., G. Martin and T. Lenormand, 2011 Fitness landscapes: An alternative theory for the dominance of mutation. Genetics 189: 923–937.

Maynard Smith, J., and J. Haigh, 1974 The hitch-hiking effect of a favourable gene. Genet. Res. 23: 23–35.

Montgomery, M. E., L. M. Woodworth, P. R. England, D. A. Briscoe and R. Frankham, 2010 Widespread selective sweeps affecting microsatellites in *Drosophila* populations adapting to captivity: implications for captive breeding programs. Biol. Cons. 143: 1842–1849.

Neher, R. A., 2013 Genetic draft, selective interference and population genetics of rapid adaptation. Ann. Rev. Ecol. Evol. Syst. 44: 195–215.

Ohta, T., 1971 Associative overdominance caused by linked detrimental mutations. Genet. Res. 18: 277–286.

Ohta, T., 1973 Effect of linkage on behaviour of mutant genes in finite populations. Theor. Pop. Biol. 4: 145–172.

Ohta, T., and C. C. Cockerham, 1974 Detrimental genes with partial selfing and effects on a neutral locus. Genet. Res. 23: 191–200.

Ohta, T., and M. Kimura, 1970 Development of associative overdominance through linkage disequilibrium in finite populations. Genet. Res. 18: 277–286.

Ohta, T., and M. Kimura, 1971 Linkage disequilibrium between two segregating nucleotide sites under steady flux of mutations in a finite population. Genetics 68: 571–580.

Palsson, S., 2001 The effects of deleterious mutations in cyclically parthenogetic organisms. J. Theor. Biol. 208: 201–214.

Pamilo, P., and S. Palsson, 1998 Associative overdominance, heterozygosity and fitness. Heredity 81: 381–389.

Pamilo, P., and S. Palsson, 1999 The effects of deleterious mutations on linked neutral variation in small populations. Genetics 153: 475–483.

Price, G. R., 1970 Selection and covariance. Nature 227: 520–521.

Robertson, A., 1962 Selection for heterozygotes in small populations. Genetics 47: 1291–1300.

Rumball, W., I. R. Franklin, R. Frankham and B. L. Sheldon, 1994 Decline in heterozygosity under full-sib and double first-cousin inbreeding in *Drosophila melanogaster*. Genetics 136: 1039–1049.

Santiago, E., and A. Caballero, 1995 Effective size of populations under selection. Genetics 139: 1013–1030.

Sved, J. A., 1968 The stability of linked systems of loci with a small population size. Genetics 59: 543–563.

Sved, J. A., 1971 Linkage disequilibrium and homozygosity of chromosome segments in finite populations. Theor. Pop. Biol. 2: 125–141.

Sved, J. A., 1972 Heterosis at the level of the chromosome and at the level of the gene. Theor. Pop. Biol. 3: 491–506.

Szulkin, M., N. Bierne and P. David, 2010 Heterozygosity fitness correlations: a time for reappraisal. Evolution 64: 1202–1217.

Wang, J. and W. G. Hill, 1999 Effect of selection against deleterious mutations on the decline in heterozygosity at neutral loci in closely inbreeding populations. Genetics 153: 1475–1489.

Wang, J., W. G. Hill, D. Charlesworth and B. Charlesworth, 1999 Dynamics of inbreeding depression due to deleterious mutations in small populations: mutation parameters and inbreeding rate. Genet Res 74: 165–178.

Weir, B. S., P. J. Avery and W. G. Hill, 1980 Effect of mating structure on variation in inbreeding coefficient. Theor. Pop. Biol. 18: 396–429.

Whitlock, M.J. and N. H. Barton, 1997 The effective size of a subdivided population. Genetics 146: 427–441.

Willams, J. L., S. J. G. Hall, M. Del Corvo, K. T. Ballingall, L. Colli et al., 2016 Inbreeding and purging at the genomic level: the Chillingham cattle reveal extensive, non-random SNP heterozygosity. Animal Genet. 47: 19–27.

